# Worldwide Soundscapes: a synthesis of passive acoustic monitoring across realms

**DOI:** 10.1101/2024.04.10.588860

**Authors:** Kevin FA Darras, Rodney Rountree, Steven Van Wilgenburg, Anna F Cord, Youfang Chen, Lijun Dong, Agnès Rocquencourt, Camille Desjonquères, Patrick Mauritz Diaz, Tzu-Hao Lin, Amandine Gasc, Sarah Marley, Marcus Salton, Laura Schillé, Paul Jacobus Wensveen, Shih-Hung Wu, Orlando Acevedo-Charry, Matyáš Adam, Jacopo Aguzzi, Irmak Akoglu, M Clara P Amorim, Michel André, Alexandre Antonelli, Leandro Aparecido Do Nascimento, Giulliana Appel, Stephanie Archer, Christos Astaras, Andrey Atemasov, Jamieson Atkinson, Joël Attia, Emanuel Baltag, Luc Barbaro, Fritjof Basan, Carly Batist, Adriá López Baucells, Julio Ernesto Baumgarten, Just T Bayle Sempere, Kristen Bellisario, Asaf Ben David, Oded Berger-Tal, Matthew G Betts, Iqbal Bhalla, Thiago Bicudo, Marta Bolgan, Sara Bombaci, Martin Boullhesen, Tom Bradfer-Lawrence, Robert A Briers, Michal Budka, Katie Burchard, Giuseppa Buscaino, Alice Calvente, Maria Ceraulo, Maite Cerezo-Araujo, Gunnar Cerwén, Maria Chistopolova, Christopher W Clark, Kieran D Cox, Benjamin Cretois, Chapin Czarnecki, Luis P da Silva, Wigna da Silva, Laurence H De Clippele, David de la Haye, Ana Silvia de Oliveira Tissiani, Devin de Zwaan, M. Eugenia Degano, Joaquin del Rio, Christian Devenish, Ricardo Díaz-Delgado, Pedro Diniz, Dorgival Diógenes Oliveira-Júnior, Thiago Dorigo, Saskia Dröge, Marina Duarte, Adam Duarte, Kerry Dunleavy, Robert Dziak, Simon Elise, Hiroto Enari, Haruka S Enari, Florence Erbs, Britas Klemens Eriksson, Pınar Ertör-Akyazi, Nina Ferrari, Luane Ferreira, Abram B Fleishman, Paulo Fonseca, Bárbara Freitas, Nick Friedman, Jérémy SP Froidevaux, Svetlana Gogoleva, Maria Isabel Gonçalves, Carolina Gonzaga, José Miguel González Correa, Eben Goodale, Benjamin Gottesman, Ingo Grass, Jack Greenhalgh, Jocelyn Gregoire, Jonas Hagge, William Halliday, Antonia Hammer, Tara Hanf-Dressler, Sylvain Haupert, Samara Haver, Daniel Hending, Jose Hernandez-Blanco, Thomas Hiller, Joe Chun-Chia Huang, Katie Lois Hutchinson, Carole Hyacinthe, Janet Jackson, Alain Jacot, Olaf Jahn, Francis Juanes, Ellen Kenchington, Sebastian Kepfer-Rojas, Justin Kitzes, Tharaka Kusuminda, Yael Lehnardt, Jialin Lei, Paula Leitman, José Leon, Deng Li, Cicero Simão Lima-Santos, Kyle John Lloyd, Audrey Looby, David López-Bosch, Tatiana Maeda, Franck Malige, Christos Mammides, Gabriel Marcacci, Matthias Markolf, Marinez Isaac Marques, Charles W Martin, Dominic A Martin, Kathy Martin, Matthew McKown, Logan JT McLeod, Oliver Metcalf, Christoph Meyer, Grzegorz Mikusinski, João Monteiro, Larissa Sayuri Moreira Sugai, Dave Morris, Sandra Müller, Sebastian Eduardo Muñoz-Duque, Kelsie A Murchy, Ivan Nagelkerken, Maria Mas Navarro, Rym Nouioua, Carolina Ocampo-Ariza, Julian D Olden, Steffen Oppel, Anna N Osiecka, Elena Papale, Miles Parsons, Julie Patris, João Pedro Marques, Filipa Isabel Pereira Samarra, Cristian Pérez-Granados, Liliana Piatti, Mauro Pichorim, Thiago Pinheiro, Jean-Nicolas Pradervand, John Quinn, Bernardo Quintella, Craig Radford, Xavier Raick, Ana Rainho, Emiliano Ramalho, Sylvie Rétaux, Laura K Reynolds, Klaus Riede, Talen Rimmer, Noelia Rios, Ricardo Rocha, Luciana Rocha, Paul Roe, Samuel RP-J Ross, Carolyn M Rosten, Carlos Salustio-Gomes, Philip Samartzis, José Santos, Kevin Scharffenberg, Renée P Schoeman, Karl-Ludwig Schuchmann, Esther Sebastián-González, Sebastian Seibold, Sarab Sethi, Fannie Shabangu, Taylor Shaw, Xiaoli Shen, David Singer, Ana Sirovic, Brittnie Spriel, Jenni Stanley, Valeria da Cunha Tavares, Karolin Thomisch, Jianfeng Tong, Laura Torrent, Juan Traba, Junior A Tremblay, Leonardo Trevelin, Sunny Tseng, Mao-Ning Tuanmu, Théophile Turco, Marisol Valverde, Ben Vernasco, Manuel Vieira, Raiane Vital da Paz, Matthew Ward, Maryann Watson, Matthew Weldy, Julia Wiel, Jacob Willie, Heather Wood, Jinshan Xu, Wenyi Zhou, Songhai Li, Renata Sousa-Lima, Thomas Cherico Wanger

## Abstract

**Aim:** The urgency for remote, reliable, and scalable biodiversity monitoring amidst mounting human pressures on ecosystems has sparked worldwide interest in Passive Acoustic Monitoring (PAM), which can track life underwater and on land. However, we lack a unified methodology to report this sampling effort and a comprehensive overview of PAM coverage to gauge its potential as a global research and monitoring tool. To remediate this, we created the Worldwide Soundscapes project, a collaborative network and growing database comprising metadata from 409 datasets across all realms (terrestrial, marine, freshwater, and subterranean).

**Location:** Worldwide, 12 200 sites, all ecosystems.

**Time period:** 1991 to present.

**Major taxa studied:** All soniferous taxa.

**Methods:** We synthesise sampling coverage across spatial, temporal, and ecological scales using metadata describing sampling locations, deployment schedules, focal taxa, and recording parameters. We explore global trends in biological, anthropogenic, and geophysical sounds based on 168 recordings from twelve ecosystems across all realms.

**Results:** Terrestrial sampling is spatially denser (45 sites/Mkm^2^) than aquatic sampling (0.3 and 1.8 sites/Mkm^2^ in oceans and fresh water) with only two subterranean datasets. Although diel and lunar cycles are well-covered in all realms, only marine datasets (56%) comprehensively sample all seasons. Across twelve ecosystems, biological sounds show contrasting diel patterns, decline with distance from the equator and negatively correlate with anthropogenic sounds.

**Main conclusions:** PAM can inform macroecology studies as well as global conservation and phenology syntheses, but representation can be improved by expanding terrestrial taxonomic scope, sampling coverage in the high seas and subterranean ecosystems, and spatio-temporal replication in freshwater habitats. Overall, this global PAM network holds promise to support global biodiversity research and monitoring efforts.

## Introduction

Sounds permeate all realms on Earth — terrestrial, freshwater, marine, and subterranean^1^. Passive Acoustic Monitoring (PAM) captures soundscapes that document soniferous (i.e., sound-producing) organisms, human activities, and some geophysical events (i.e., biophony, anthropophony, and geophony, respectively). In ecoacoustics and soundscape ecology^2,3^, PAM can measure levels and impacts of global change (e.g., climate change, urbanization, deep-sea mining)^4–6^; monitor ecosystem health, recovery, and restoration^7–9^; assess human-environment interactions (e.g., public health, cultural ecosystem services)^10,11^; and guide environmental management and conservation policies (e.g., protected areas, landscape planning)^12,13^.

Despite the wide-ranging and increasing soundscape sampling effort^14,15^, its global distribution across realms remains undescribed. Soundscape-recording communities are currently only networked within realms, often differing methodologically. Previous reviews focused on single realms and were either systematic^14–17^ or non-systematic qualitative^18,19^ reviews that cannot describe data trends. The systematic reviews did not quantitatively address taxonomic focus in marine studies, nor ecosystemic terrestrial coverage, or spatio-temporal sampling distribution in the freshwater realm, while the subterranean realm has not been reviewed yet. Notably, all reviews so far used only published data and involved only a small part of their respective communities. Marine scientist networks using PAM exist^20^, but the freshwater community is nascent, and the terrestrial community is fragmented among single taxa. Methodological differences are also striking: acoustic calibration and sound propagation modelling are advanced in aquatic studies^21^ but seldom considered in terrestrial ones (but see^22,23^); artificial intelligence can identify increasing numbers of species on land^24^, whereas most aquatic sounds are still challenging to identify^25,26^.

Overall, building a united global PAM network can increase knowledge transfer resulting in more efficient and consistent methods, analyses, and cross-system syntheses^14^. Cross-realm PAM studies can yield new theoretical answers^27^ and applied solutions: organisms’ sound durations follow a common distribution across multiple realms^28^; soundscapes track terrestrial and marine resilience to disturbance regimes^29^. Transnational sampling could form the basis for comprehensive soniferous biodiversity monitoring, just as community-initiated telemetry databases^30^ and collaborative camera trap surveys^31^ have advanced entire research fields. A global PAM network could complement existing biodiversity-monitoring networks, establish historical biodiversity baselines, support systematic long-term and large-scale monitoring, and connect with the public through citizen science. Such information is critical to inform global biodiversity policies^32^ such as the Kunming-Montreal Global Biodiversity Framework.

We present the “Worldwide Soundscapes” project, the first global PAM meta-database and network (https://ecosound-web.de/ecosound_web/collection/index/106). We used it to quantify the known state of PAM efforts, highlight apparent sampling gaps and biases, illustrate the potential of cross-realm PAM syntheses for research, and federate PAM users. The project currently comprises 351 active contributors who collated metadata from 409 passively-recorded, stationary, replicated soundscape datasets. Metadata describe the exact spatio-temporal coverage, sampled ecosystems (International Union for Conservation of Nature Global Ecosystem Typology: IUCN GET), transmission medium (air, water, or soil), focal taxa (IUCN Red list), recording settings, as well as data and publication availability. We inferred coverage within administrative (Global ADMinistrative Database: GADM; International Hydrographic Organisation: IHO) and protected areas (World Database on Protected Areas: WDPA) from geographic locations. We selected recordings referenced in the meta-database to quantify soundscape components (biophony, anthropophony, and geophony) across twelve ecosystems from all realms. We showcase their relevance to exemplary macroecology, conservation biology, and phenology research questions and identify opportunities to advance the global PAM network. The publicly-accessible meta-database^33^ continues to grow to enhance accessibility of data and remains open for metadata contributions, facilitating future syntheses.

## Methods

### Database construction

The database construction started in August 2021 within the frame of the Worldwide Soundscapes project using collaborative, peer-driven metadata collation^33^. It represents the current state of knowledge of PAM within our network. We additionally conducted focal publication searches to plug coverage gaps by inviting the respective corresponding authors. We posted the call for contributors on specialised ecoacoustics platforms and social media, and keep the project open for any contributor owning suitable soundscape recordings. We communicated by emails in English, Spanish, French, Portuguese, German, Russian, and Chinese to federate users in our network. Out of 592 contacted contributors potentially involved in PAM, 63% already provided metadata, 23% have not yet responded, 12% are pending, and 2% have declined. We included metadata from larger groups such as the Silent Cities project, Ocean Networks Canada, and the Australian Acoustic Observatory. Primary contributors provided the metadata and bear the responsibility of their accuracy. Research assistants checked the coherence of the metadata input (beyond automated data format checks) and primary contributors cross-validated metadata displayed as maps and graphical timelines. The database information page and content is integrated in our online collaborative ecoacoustics platform ecoSound-web^34^ that also hosts the annotated soundscape recordings of the case study (https://ecosound-web.de/ecosound_web/collection/show/49).

Soundscape recording datasets were required to meet four criteria: 1) stationary — mobile recorders have variable spatial assignments, thus we excluded recordings from cars, transect walks, or towed deployments; 2) passive — obtained from unattended recorders; 3) ambient — omni-directional, non-triggered recordings, under non-experimental conditions; 4) spatially or temporally replicated (Fig. 1) — as to disentangle spatial and temporal effects from other soundscape determinants. Datasets were defined as spatially replicated when several sites were sampled simultaneously, and temporally replicated when a site was sampled over multiple days at the same time of day. Sampling sites and days were our elemental units for defining replication, however, in other contexts, spatial replicates may for instance be required to be in the same habitat, and temporal replicates could be defined across multiple full moon nights. Taken together, our requirements homogenise the dataset to enable general, unified statistical analyses across datasets for future syntheses.

**Figure 1:**
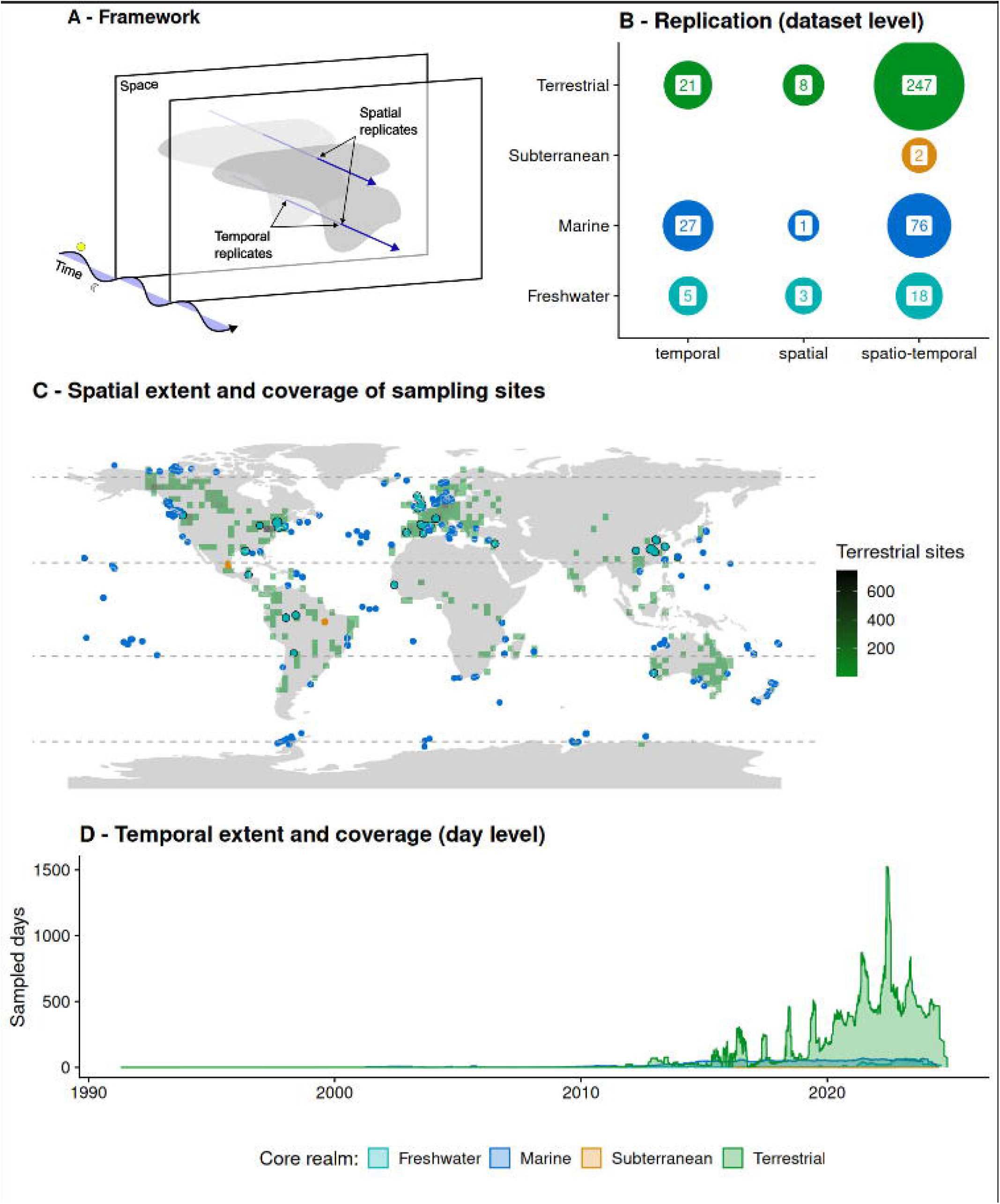
Overview of Worldwide Soundscapes meta-database. A) Framework used to define spatial and temporal replicates. B) Number of datasets in each core realm for the different replication levels. C) Spatial extent and coverage, based on sampling sites, split by core realm. Due to their higher representation and to avoid overlapping site clusters, terrestrial site densities were plotted on a 3 degree resolution raster (Interactive map: https://ecosound-web.de/ecosound_web/collection/index/106). D) Temporal extent and coverage, based on recorded days, split by core realm. An enlarged version of panel D without terrestrial sites can be found in Fig S3. For panels C and D, sites from transitional realms were assigned to their parent core realm.

**Figure 2:**
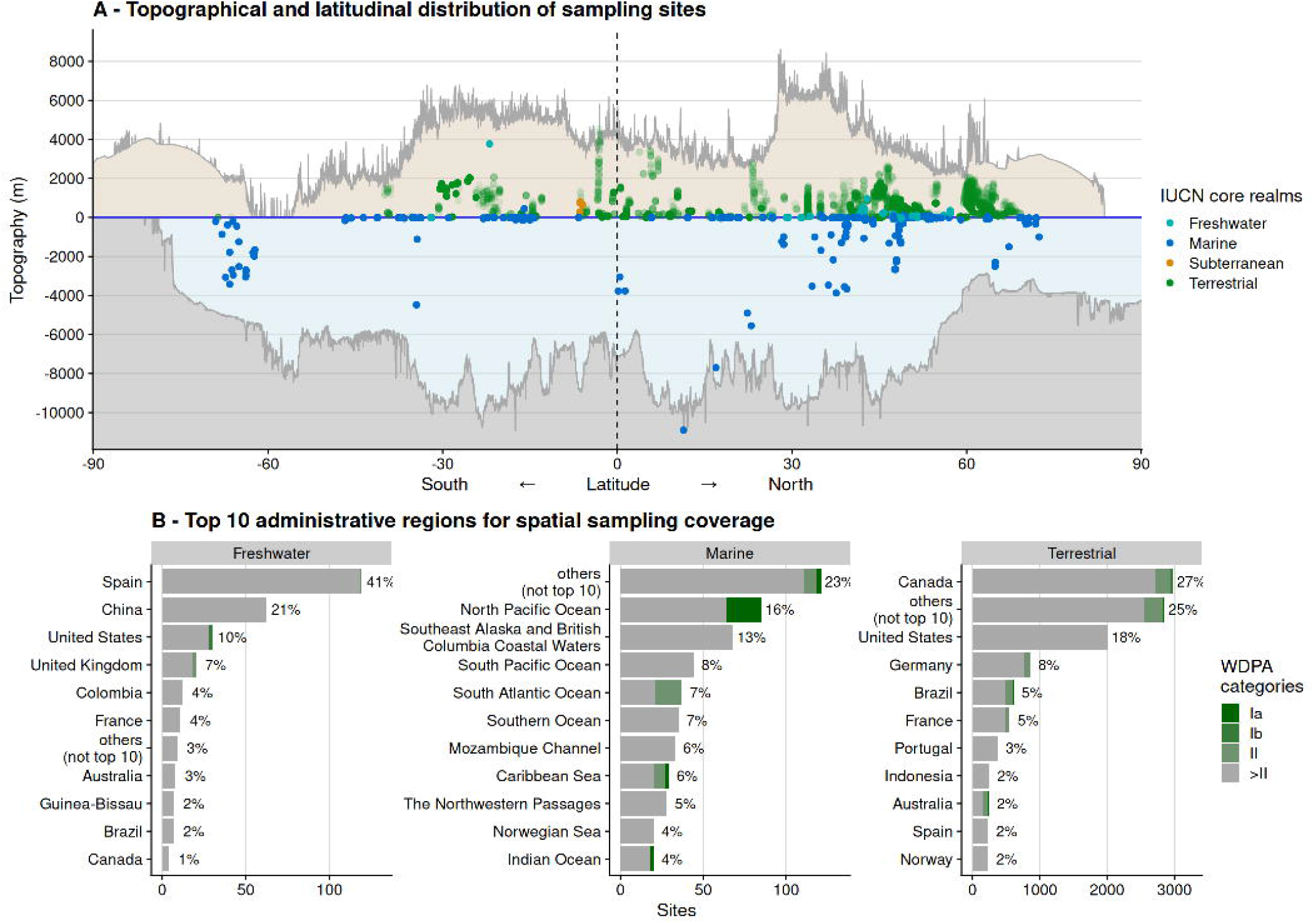
Spatial distribution of sampling sites. A) Latitudinal and topographic distribution of sampling sites across core realms. Due to their higher representation and to avoid overlapping site clusters, terrestrial sites are shown with transparency. The minimum (deepest seafloor) and maximum (highest elevation of land or sea level) topographical limits (dark grey lines) are shown against latitude, based on General Bathymetric Chart of the Oceans data36. Minimum topography above the sea level and maximum topography under the sea level were set to zero as the sea level represents the minimum and maximum in these cases. B) Number of sampling sites within different administrative regions (GADM level 0 and IHO sea areas), split by core realm, across WDPA categories (Ia: strict nature reserve; Ib: wilderness area, II: national park). The areas that do not belong to the top 10 in terms of datasets have been aggregated under “others”. Six subterranean sites in Brazil are not shown. Sites from transitional realms were assigned to their parent core realm.

### Time and space

Soundscapes result from geophysical phenomena as well as wildlife and human activities that are broadly determined by solar and lunar cycles, and geographical positions on the planet relative to the poles or equator, or the land and water surface. We defined and calculated spatial coverage as the number of sites, and spatial density as the number of sites per the containing realms’ areal extent. Alternatively, spatial extent could have been defined as the area bounded by sites, but calculating extents on the world sphere is conceptually challenging for large extents. Further, the sampling area covered at each site is generally unknown: they are rarely measured in terrestrial sites but sometimes simulated in marine environments^35^. Detection spaces vary with sound source intensity, frequency, directivity; recording medium temperature, currents, pressure; habitat structure; and ambient sound level. Underwater, detection spaces are greater due to the higher density of the recording medium. Our measure of spatial sampling density using points per area is thus provisional. Temporal extent was defined and calculated as the time range from the start of the first to the end of the last recording, temporal coverage as the time sampled per site, and temporal density as the proportion of time sampled within the temporal extent.

We quantified latitudinal and topographical distribution by collecting coordinates, altitude, and depth data for each site. Topography values on land were either provided by the contributors or filled in using General Bathymetric Chart of the Oceans surface elevation data^36^. For underwater sites, depth values below the water surface were provided by the contributors, and for subterranean sites, depth below the land surface was recorded. We assigned sampling sites to administrative areas (GADM divisions for freshwater and land, IHO for sea areas) and extracted their WDPA category. Sites’ climates were geographically classified into tropical (between −23.5° and 23.5° latitude), polar (below −66.5° or above 66.5° latitude), and temperate (between polar and tropical regions).

Primary contributors coded their exact recording times for each site using deployment start and end dates and times and operation modes (or combinations thereof). Continuous operations lasted from the deployment start to its end; scheduled operations had daily start and end times; periodical operations used duty cycles. A temporal framework was devised to quantify sampling coverage in three solar and lunar cycles using the timing of sound recordings relative to time events (Fig. 3). Seasonal coverage was inferred only for temperate sites, by splitting the year into four meteorological seasons (winter: December-February, spring: March-May, summer: June-August, fall: September-November, reversed for Southern latitude sites). The daily cycle was split into four diel windows delimiting dawn (from astronomical dawn start at -18° solar altitude until 18° solar altitude), day, dusk (from 18° solar altitude to astronomical dusk end at -18° solar altitude), and night. The lunar illumination cycle was split into two time windows centred on the full and new moon phases. Thus, extrema and ecotones in the temporal cycles define time windows, and in temperate zones, equinoxes roughly correspond to thermal ecotones. Seasonal cycles in tropical and polar regions arising from precipitation patterns were not considered in this analysis.

**Figure 3:**
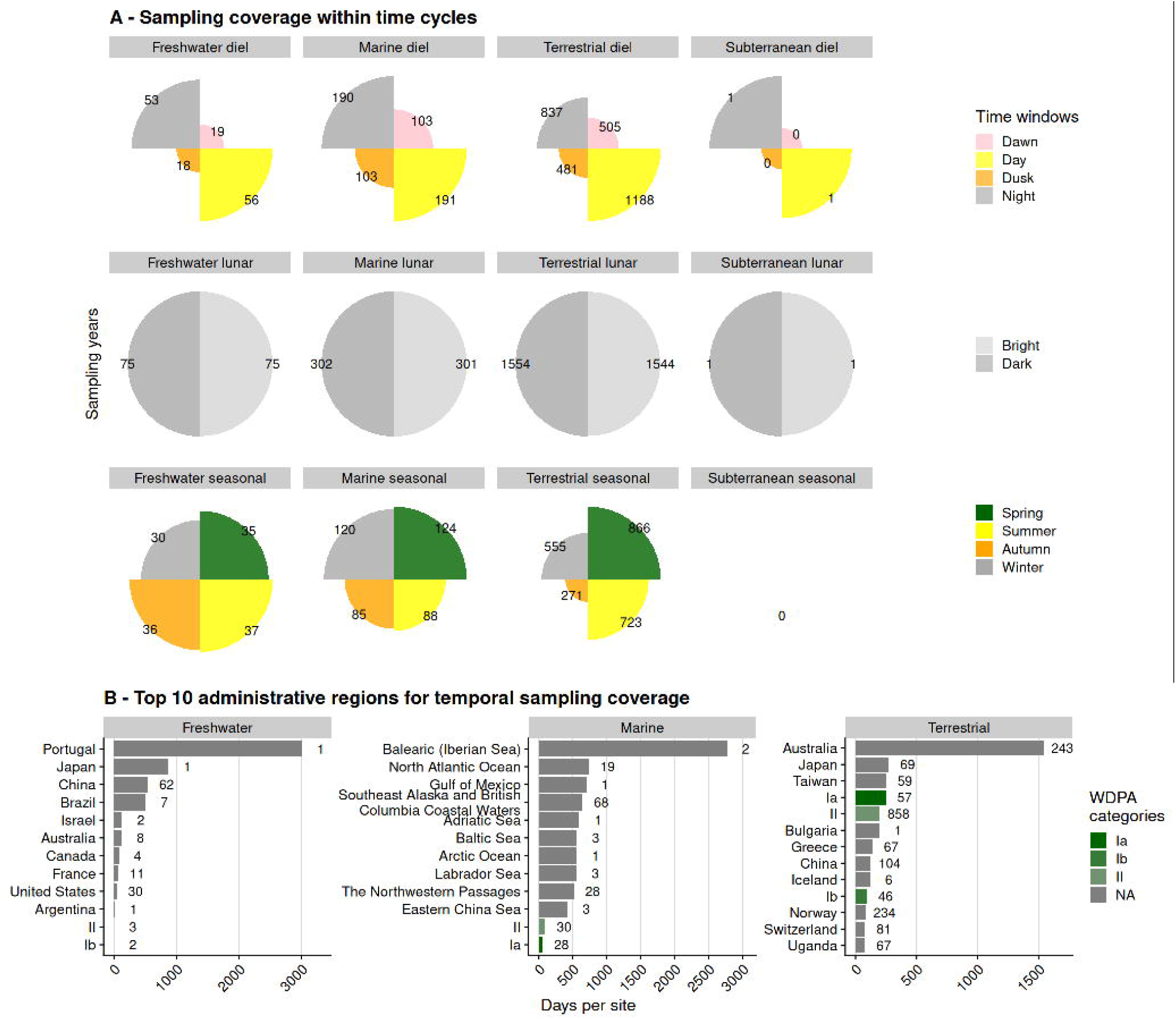
Temporal sampling distribution. A) Temporal sampling coverage across solar and lunar cycles for the three core realms. Cycles consist of solar (daily and seasonal) and lunar time cycles (lunar phase), subdivided in time windows. Seasons were only analysed in temperate regions. Sampling coverage is represented with sampling days in number labels. B) Mean number of sampling days per site within different administrative regions (GADM level 0, IHO areas, WDPA categories), split by core realm. WDPA categories (Ia: strict nature reserve; Ib: wilderness area, II: national park) are shown separately. Six subterranean sites in Brazil are not shown. Numbers to the right of bars indicate the number of sites the means were calculated from. Sites from transitional realms were assigned to their parent core realm.

### Ecological characterisation

We assigned individual sampling locations to ecosystem types following the IUCN GET (https://global-ecosystems.org). Sites were assigned hierarchically to realms (core or transitional ones, which represent the interface between core realms), biomes, and functional groups. We calculated major occurrence areas of all functional groups based on ecosystem maps^37^ to quantify spatio-temporal extent, coverage, and sampling density within realms and biomes. Notably, only the result section “Sampling in ecosystems’’ separates results among core and transitional realms. For the spatial and temporal result sections, sites were assigned to their “parent” realm (i.e., the first mentioned realm in the compound transitional realm names). Deployments were linked to IUCN Red List taxa (class, order, family, or genus) when studies were designed for monitoring these taxa, and other deployments could be collected without taxonomic focus.

### Acoustic frequency ranges

Sound recordings store a representation of the original soundscape, so we needed to determine their spectral scope. Microphones (including hydrophones and geophones) have variable frequency responses, usually declining with frequencies above the human-audible range. Additionally, digital recorders restrict the spectral scope of the recording with the sampling frequency. Contributors provided audio parameters for their deployments: sampling rate, high-pass filters, microphone and recorder models.

### Soundscape case studies

To illustrate how the database can be used for macroecology, conservation biology, and phenology analyses, we selected 168 recordings across a variety of topographical, latitudinal, and anthropisation conditions — all fundamental gradients of assembly filters in both terrestrial and marine realms^1^ — belonging to 12 IUCN GET functional groups. We aimed for four spatial replicates within the same functional group, with 10-minute audible sound recordings (at least 44.1 kHz sampling frequency) starting at sunrise, solar noon, sunset, and solar midnight from the same date during the biologically active season. The available data did not always allow this (Table S2). We acknowledge that this targeted, non-systematic selection is not statistically representative of global patterns but rather illustrative of the database potential.

Each soundscape recording was analysed with respect to the three fundamental soundscape components: biophony, anthropophony, and geophony. Soundscape recordings were uploaded to ecoSound-web^34^ for annotation (https://ecosound-web.de/ecosound_web/collection/show/49): KD listened to them while visually inspecting their spectrograms (Fast Fourier Transform window size of 1 024) at a density of 1 116 pixels per 10 minutes. Annotations were rectangular boxes with defined coordinates in the time and frequency dimensions on the spectrogram, bounding only the annotated sound. Annotations of different soundscape components could overlap if they were simultaneously visible or audible. The total expanse of annotations of the same soundscape component was computed, excluding overlaps. Soundscape components above 22.05 kHz and sounds caused by microphone or recorder self-noise were excluded. All annotations were reviewed by the recordists using the peer-review mode on ecoSound-web. Acoustic space occupancy for each soundscape component in each recording was calculated as the proportion of the sampled spectro-temporal space^38^ (i.e., total annotation area divided by total area of spectrogram; range: 0-1).

We asked whether biophony occupancy increases with proximity to the equator, whether biophony was negatively correlated with anthropophony, and whether phenology patterns differ across realms. Statistical models predicting biophony occupancy were Bayesian beta regression models (4 chains of 1000 sampling iterations with 1000 warmup iterations) fitted with the R package brms^39^. Models converged as determined by trace plots and R hat values smaller than 1.1. The number of samples equaled the number of recordings (N=168). The models using latitude and anthropophony were mixed-effect models including the functional group as a random intercept. The phenology model used diel time windows and realms, as well as their interactions, as predictors.

## Results

### Summary dataset statistics

To date, 409 validated soundscape meta-datasets (hereafter “datasets”) have been registered in our database from across the globe, dating back to 1991 (Fig. 1D). A dataset gathers a team’s metadata on a study or project. Based on the IUCN GET definition of four ‘core’ realms, our database includes 277, 104, 26, and 2 validated datasets from the terrestrial, marine, freshwater, and subterranean realms, respectively. The transmission medium was air for terrestrial datasets (except for soil in one dataset), mostly water for aquatic datasets (eight above-water datasets mostly in freshwater realm). In the subterranean realm, one dataset did aerial recordings, while the other did underwater recordings. The majority of datasets (84%) include both spatial and temporal replicates (Fig. 1B). Few datasets have openly-accessible recordings (10-12% excluding subterranean realm, Fig. S1). Presently, few terrestrial and freshwater datasets (27% and 19%, respectively) are associated with DOI-referenced publications in contrast to marine datasets (44%, excluding subterranean realm) (Fig. S1).

### Spatial sampling coverage and density

The database contains 12 220 sampling sites, including 147 polar, 9 194 temperate, and 2 859 tropical sites (Fig 1C). On land, 11 229 sites are located within 85 (out of 263) GADM level 0 areas (i.e., countries, Fig. 2B), primarily in the Northern Hemisphere (Table S1). Administratively, most terrestrial sites occur in Canada (27%), followed by the United States (18%), but a significant proportion is globally widely distributed (25% do not belong to the top 10 GADM areas). Few terrestrial sites (8%) are located in WDPA category Ia, Ib, or II areas, corresponding to the highest protection levels. Our database currently lacks data from vast areas in Russia, Greenland, the Antarctic, North Africa, and Central Asia. Site elevations range from sea level up to 4 548 m (Fig. 2A), but mountains above 4 000 m in the Northern Hemisphere, and above 2 000 m in the Southern Hemisphere (except for the Kilimanjaro), as well as the Transantarctic Mountains, are currently not represented in the data. At sea, 634 sites are located within 35 (out of 101) IHO sea areas. Administratively, most marine sites are widespread among IHO areas (23% do not belong to the top 10 IHO areas), but the North Pacific Ocean and the Southeast Alaskan and British Columbian coastal waters contain large parts of marine sites (16% and 13%, respectively). Many sites are situated in WDPA high-protection category Ia and II areas (12%). Our database currently lacks datasets from Arctic waters off Eurasia, and Southeast Asian coastal areas. Sampling sites span ocean depths from sea water surface to depths of 10 090 m, but tropical bathypelagic and Southern benthic areas are poorly represented (Table S2). Few GADM areas (16) are represented in the 324 freshwater sites. Spain holds most freshwater sites (56%), followed by China (21%), and few freshwater sites (2%) are in WDPA category II or Ib areas.

Freshwater bodies are sampled at elevations from sea level up to 3 770 m and were sampled up to 25 m depth. Mountain freshwater bodies and those in Africa, Asia, and Oceania are currently poorly represented in our database (only one site at 3770 m). The database contains 13 subterranean sites situated in Brazil and Mexico between 277 and 810 m elevation, and up to a depth of 1 546 m (horizontal projection).

### Temporal sampling extent, coverage, and density

We compare sampling coverage (in years sampled by recordings, summed over sites) across time windows of the diel cycle, the lunar phases, and the seasons (Fig. 3A). Dawn and dusk diel windows are shorter than day and night diel windows for most locations and correspondingly less intensively sampled. The lunar phase cycle is evenly covered across realms and comprehensively covered within datasets (Fig. S2). In the terrestrial realm, daytime coverage surpasses nighttime coverage (606 vs. 395 years), while 74% of datasets sampled all diel time windows. Terrestrial temperate datasets mostly sampled spring (411 years, 32%) and summer (559 years, 44%) while 26% sampled all seasons. Terrestrial temporal coverage per site is highest in Australia (1541 days). In the marine realm, diel coverage is even, as 89% of marine datasets sampled all diel time windows. Marine temperate datasets have high and similar coverage for winter and spring (117 and 122 years, combined 59% of seasonal coverage) and 64% cover the full seasonal cycle. Marine temporal coverage per site is highest in the Balearic (2771 days). In the freshwater realm, temporal coverage among diel time windows is even and 81% of freshwater temperate datasets sampled all diel time windows. Similarly, 13% sampled all seasons. Freshwater temporal coverage per site is highest in Portugal (3 009 days). The subterranean tropical sites primarily covered the nighttime.

### Sampling in ecosystems

Our database includes 83 of the 110 IUCN GET functional groups and all biomes except anthropogenic shorelines, the subterranean tidal biome, anthropogenic subterranean voids, and anthropogenic subterranean freshwaters (Table S2). The terrestrial realm has the third-largest extent and the highest spatial sampling density among realms (45 sites per Mkm^2^ over entire temporal extent), but temporal coverage is comparatively low (30% sampled out of 203 days of mean extent per site). The most commonly sampled terrestrial biome is the temperate-boreal forests and woodlands biome (56% of sites). The marine realm is the most extensive but spatial sampling density is the lowest (0.3 sites per Mkm^2^), while temporal sampling coverage is the highest among all realms (66% out of 379 days sampled). The most commonly sampled marine biomes are the marine shelf and pelagic ocean waters (56% and 33% of sites respectively). The freshwater realm has low spatial sampling densities (1.8 sites per Mkm^2^) and high temporal sampling densities (67% out of 238 days sampled). Lakes are the most commonly sampled freshwater biome (48% of sites). The terrestrial-freshwater realm, representing 81% of the area of non-subterranean realms, has the third-highest spatial sampling density (7.9 sites per Mkm^2^) and similar temporal sampling density to the terrestrial realm (18% out of 233 days sampled). The marine-freshwater-terrestrial realm (including 26 sites in coastal river deltas, saltmarshes, and intertidal forests) has the largest temporal extent (465 days per site). The subterranean realm, though second-largest, includes seven tropical sites (all in tropical aerobic caves) sampled with a low temporal coverage (1% out of 129 days sampled), and the subterranean-freshwater realm includes five sites in underground streams and pools, sampled with a high temporal coverage (17% out of 507 days sampled).

### Target taxa and frequency ranges

Most marine datasets do not target specific taxa (66%) and use wide frequency ranges from 0.01 to 31 kHz (mean bounds of frequency ranges across datasets, Fig. 4B). Marine datasets that focus on single taxa comprise fish (12%, 0.002 - 20 kHz) and cetaceans (6%, 0.006 - 7 kHz). Similarly, most freshwater datasets are taxonomically unspecific (56%) and cover frequencies from 1 to 29 kHz. Some datasets (14%) focus on ray-finned fish, covering frequencies from 0.001 to 23 kHz. In contrast, terrestrial datasets mostly target single taxa and have narrow frequency ranges. Bird-focused datasets are most common (44%), spanning frequencies from 0.06 to 21 kHz, while bat-focused datasets are next (12%) and range from 5 to 139 kHz. Taxonomically unspecific datasets account for 24% of terrestrial datasets, covering a broad range from 0.2 to 23 kHz. Generally, datasets targeting multiple taxa use wider frequency ranges than those targeting single taxa.

**Figure 4:**
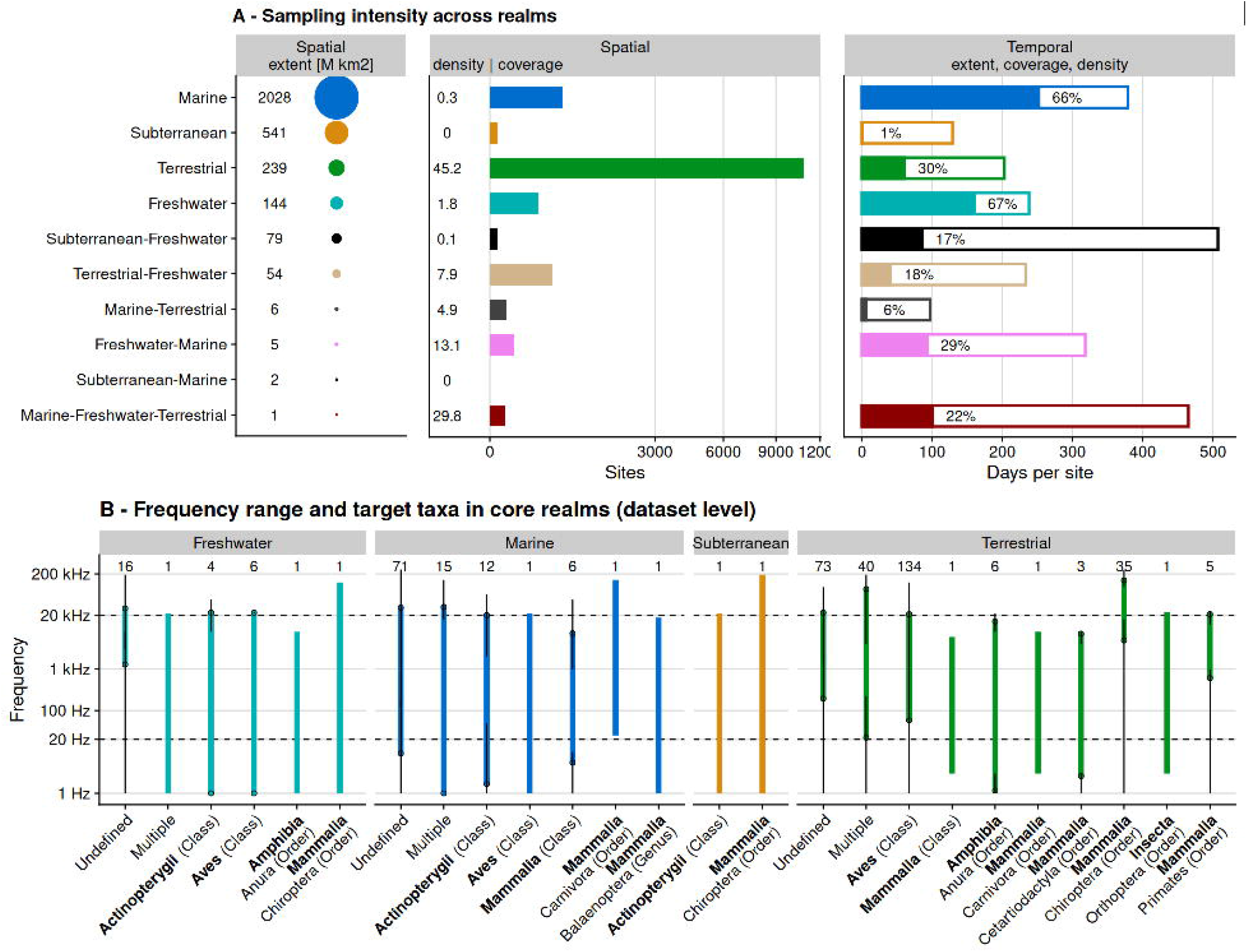
Sampling distribution across ecological scales. A) Spatial extent of realms, based on major areas according to IUCN GET (coloured disk area proportional to area); spatial sampling density (in sites per Mkm2) and coverage (in number of sites); temporal extent (mean range between first and last recording day), coverage (days sampled per site and density (proportion of days sampled per extent). B) Frequency ranges of datasets across realms (using Nyquist frequency i.e., actual recorded frequencies) for the main studied taxa. The dots at the ends of coloured lines represent means of the lowest and highest recorded frequencies, and the range between the minimum and maximum of these values are indicated with black error bars. The limits of human hearing are indicated with dashed lines. Number of datasets indicated above lines - datasets can be counted several times if they contain deployments targeting different taxa. Data from transitional realms were assigned to their parent core realm in panel B.

### Soundscape case studies

We analysed 168 recordings from twelve ecosystems (corresponding to different IUCN GET functional groups) representing the diversity of soundscapes on Earth, spanning latitudes from 69 degrees South to 67 degrees North (Table S3, Fig. 5A). Biophony dominated with an average soundscape occupation of 28% across all ecosystems. Notable examples include the photic coral reefs in Okinawa, Japan, with snapping shrimps and grunting fish choruses, and the tropical lowland rainforests in Jambi, Indonesia, with buzzing insects and echoing bird and primate songs, showing soundscape occupancy of 75% and 61%, respectively^40^. Only marine island slopes (off Sanriku, Japan) and polar outcrops (Antarctic) contained no or very little (2%) biophony, respectively. Geophony was absent in most of our soundscape samples, with the exception of high wind noise in polar outcrops (14%) and some wind in montane tropical forests and urban ecosystems (both 5%). Anthropophony occupied on average 11% of the soundscapes. Cities (Jambi, Indonesia; Montreal, Canada) exhibited the highest anthropophony (43%) with prevalent engine noise and human voices, while deep-sea mining and vessel communication signals caused high anthropophony in marine island slopes (32%)^41^. Silence occupancy was highest in bathypelagic ocean water (96%) and polar outcrops (82%).

**Figure 5:**
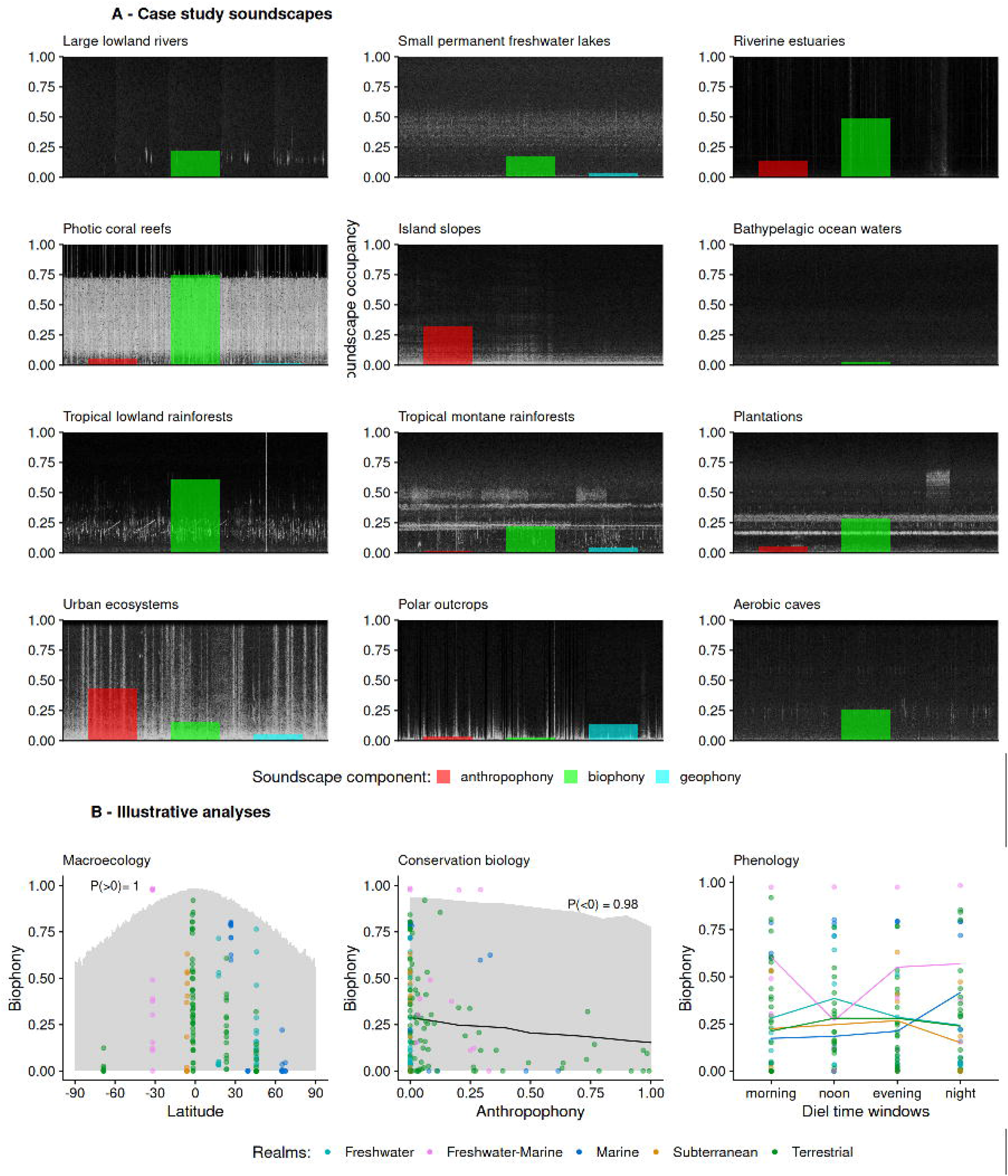
Soundscape components analysis. A) Mean acoustic space occupancy of soundscape components (biophony, geophony, anthropophony) as calculated from annotations for twelve selected ecosystems, measured in proportion of spectro-temporal space used, over 168 recordings covering the time windows of the diel cycle. Annotated recordings are accessible at https://ecosound-web.de/ecosound_web/collection/show/49. Sample spectrograms are shown in the background. B) Soundscape component occupancy data illustrate three research questions linked to macroecology (i.e., biophony with distance from equator relationship), conservation biology trade-offs (i.e., biophony with anthropophony relationship); and phenological trends (i.e., mean biophony along diel time windows for each realm). Gray ribbons indicate 95% credible intervals and numbers indicate probabilities of positive or negative relationships.

The selected soundscapes reveal greater biological activity closer to the equator, a negative relationship between biophony and anthropophony, and variable phenology patterns of soniferous organisms over the diel cycle (Fig. 5). We detected a negative correlation of biophony occupancy with increasing distance from the equator (P_negative_=1) and with anthropophony occupancy (P_negative_=0.98). The phenology model predicted biophony occupancy values for each diel time window and realm (Fig. 5B), revealing similar phenology for the terrestrial and marine realm, and opposed freshwater and freshwater-marine realm phenology.

## Discussion

The “Worldwide Soundscapes” project has — to our knowledge — assembled the first global meta-database of PAM datasets across realms. We analysed its current content to quantify sampling extent, coverage, and density across spatiotemporal and ecological scales. We analysed soundscapes from twelve ecosystems to investigate macroecological, conservation biology, and phenological trends. The database remains open for contributions and can be openly accessed to source datasets and initiate collaborative studies^33^. Next, we discuss the state and potential of PAM globally (Table 1). While we focuse on large-scale research questions here, the database can also be used to address more focused questions, for instance using the finely-resolved taxonomic information that we recorded, or for within-realm syntheses.

**Table 1:**
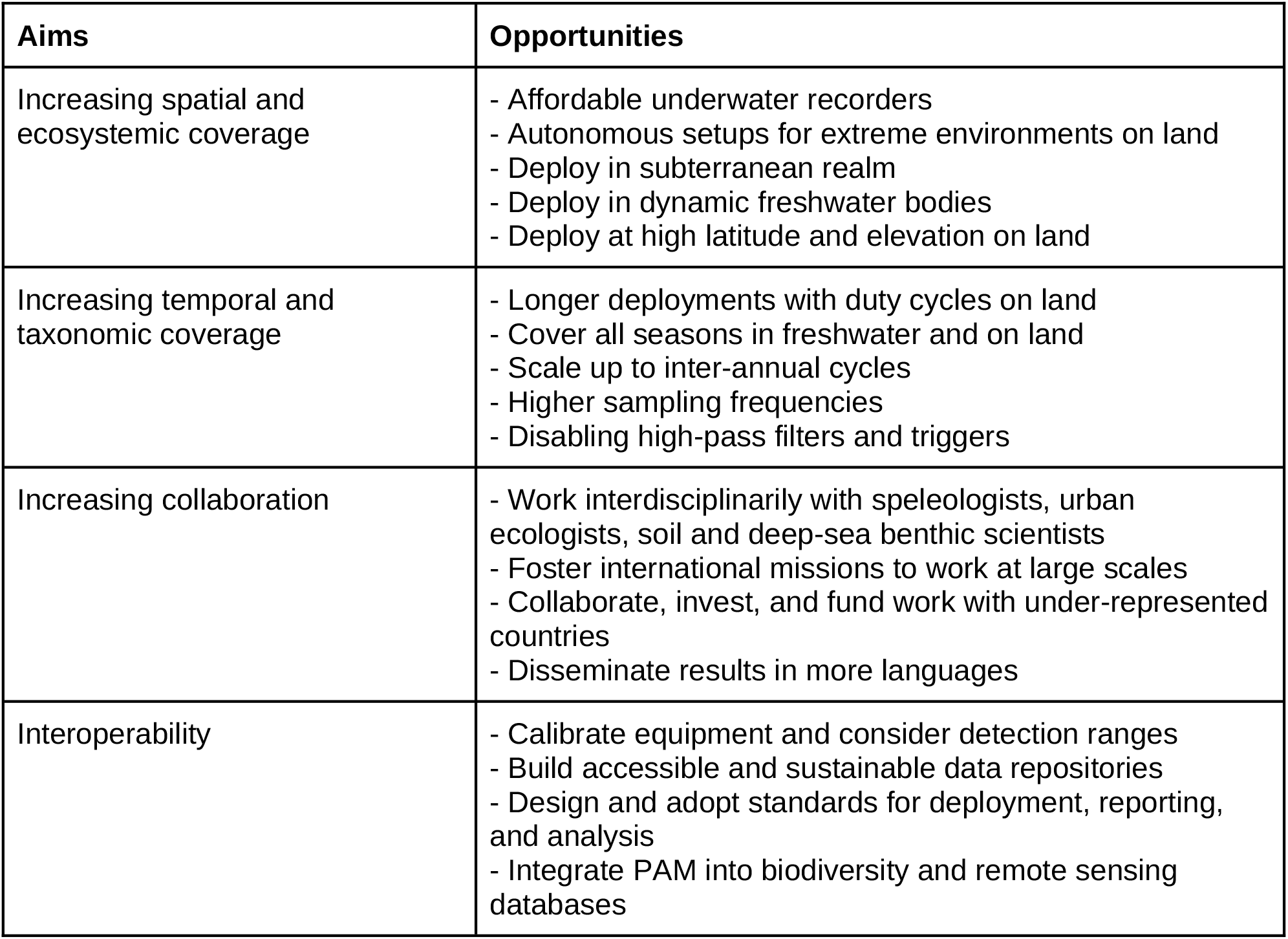
An agenda towards a global PAM coverage and network.

| Aims | Opportunities |
| --- | --- |
| Increasing spatial and ecosystemic coverage | - Affordable underwater recorders<br>- Autonomous setups for extreme environments on land<br>- Deploy in subterranean realm<br>- Deploy in dynamic freshwater bodies<br>- Deploy at high latitude and elevation on land |
| Increasing temporal and taxonomic coverage | - Longer deployments with duty cycles on land<br>- Cover all seasons in freshwater and on land<br>- Scale up to inter-annual cycles<br>- Higher sampling frequencies<br>- Disabling high-pass filters and triggers |
| Increasing collaboration | - Work interdisciplinarily with speleologists, urban ecologists, soil and deep-sea benthic scientists<br>- Foster international missions to work at large scales<br>- Collaborate, invest, and fund work with under-represented countries<br>- Disseminate results in more languages |
| Interoperability | - Calibrate equipment and consider detection ranges<br>- Build accessible and sustainable data repositories<br>- Design and adopt standards for deployment, reporting, and analysis<br>- Integrate PAM into biodiversity and remote sensing databases |

Our results likely represent global PAM trends, even though they may be biased by the project contributors’ background. Our terrestrial spatial coverage is similar to a recent systematic review^14^. Our database gaps in North Africa and Northeastern Europe correspond with the paucity of bioacoustic datasets for these regions in the Xeno-Canto bioacoustic repository^42^. Our database comprises 634 marine sampling locations, while a recent systematic review compiled 991^15^ from published data. Although the latter review’s locations are represented at the dataset level (which can comprise several sites), most overlap with our finer site-level locations (Fig S4). Marine tropical waters that are under-represented in our database reflect gaps found in the International Quiet Ocean Experiment network coverage^43^. Our marine and terrestrial database’s spatial coverage is thus broadly comparable with published data, but it is more detailed as the exact sampling times and locations are known. Our database’s temporal coverage is also more finely resolved and thus not directly comparable with previous work. To our knowledge, no other spatially-explicit review of freshwater sampling or synthesis of subterranean PAM coverage exists for comparison. Finally, as our database originates from an active network of researchers, it represents the current availability of mostly as-yet unpublished data (Fig. S1).

Geographic coverage strikingly differs among realms: marine coverage is sparse but widespread; terrestrial coverage is comparatively intensive in the Americas, Australia, and Western Europe; freshwater coverage is scattered. This partly reflects the gap between high-income countries, which have the resources to carry out conservation and active ecosystem management actions; and developing countries, which paradoxically contain the majority of biodiversity hotspots (and some of the most densely populated areas).

Accessibility and technical limits in extreme environments also drive geographic patterns: high latitudes and elevations entail extremely cold temperatures that present challenges for operation and maintenance, some of which can be solved with robust power setups (solar panels, freeze-resistant batteries). Marine deployments are generally even more constrained due to costly and demanding underwater work, but some deployments reach the polar regions as water temperatures are buffered below the freezing point. Affordable underwater recorders^44^ may help to intensify sampling of marine coastal areas and close gaps in freshwater coverage. By contrast, terrestrial monitoring is often straightforward. Northern temperate areas outside of Northeastern Europe are comparatively better covered, and the tropics outside of Africa better represented. It appears that gaps in North Africa, Central Asia, and Northeastern Europe arise from unequal research means and priorities between countries. Taken together, these gaps will help to correct spatial biases^45^ and to identify high-priority, unique research areas that should be included in global assessments.

Currently, only marine studies achieve relatively even coverage of temporal cycles. Indeed, offshore deployments — especially in the deep sea — are expensive and limited duration deployments are not cost-effective^46^. Marine soundscapes fluctuate stochastically^47^, but the ocean buffers water temperatures so that animals retain a basal activity level year-round. Although lunar phases affect marine life^48–50^ and shape tidal ecosystems, we did not consider lunar tides. In the terrestrial and freshwater realms, most deployments cover the entire diel cycle but monitoring on land often focuses either on diurnal birds or on nocturnal bats, as found in the literature^14^. In contrast, although seasons also drive acoustic activity cycles on land^51,52^, spring- and summertime monitoring is disproportionately common and we lack a thorough understanding of seasonal dynamics. Terrestrial deployments in particular may be short for logistic reasons: in the cold, batteries struggle and access is harder; in arid regions, fire hazards complicate long-term deployments; recorders are generally at risk of theft; and limited equipment may cycle between sites^53^. Lunar phases — probably evenly sampled by chance — also influence land animals^54^ but should be explicitly considered in study designs. Overall, we encourage longer-duration setups with regularly-spread sampling inside temporal cycles to alleviate the higher expenses for energy and storage, as well as trade-offs with spatial coverage^55^. Global changes impact soundscapes in largely unpredictable ways through changing species distributions and phenology, necessitating higher and unbiased coverage across multiple time scales — including inter-annual ones — to successfully monitor ongoing changes^56^.

Re-use of soundscape datasets is restricted by their taxonomic focus. Admittedly, taxonomically untargeted, long, and regular deployments in oceans, coupled to large detection ranges, concurrently sample many taxa^57^. However, many soundscape recordings sample particular frequencies, often in the human-audible range^38^, although biophony ranges from infrasound to ultrasound. For instance, studies of toothed whales or bats often use triggers and high-pass filters to cope with high data storage demands by recording purely ultrasonic recordings only when signals are detected, resulting in spectrally-restricted and temporally-biased soundscape recordings. Less-studied taxa, such as anurans and insects, could effectively be co-sampled by adjusting ongoing terrestrial deployments. We encourage terrestrial researchers to maximise frequency ranges (and to broaden diel coverage — see above) to enhance interdisciplinary collaboration. These collaborations can help to mutualise resources for mitigating potentially prohibitive power, storage, transportation, and post-processing costs^22^. Emerging embedded-AI audio detectors may offer an alternative to continuous and broadband recording^58^, but whole soundscape recordings will remain essential for broader application.

In every realm, ecosystems await acoustic discovery. Except for one dataset from aerobic caves, we currently lack data from all subterranean realms (anthropogenic voids, ground streams, sea caves), while endolithic systems may be the only ecosystem for which PAM is probably irrelevant. Access is usually challenging or restricted for non-specialists, but subterranean biodiversity shows high spatial turnover^59^. Freshwater data were less rare, and several datasets with unreplicated sampling could not be included. Temporary, dynamic water bodies (seasonal, episodic, and ephemeral ecosystems), although prevalent^60^, are not yet well-studied (Table S2) although most are accessible from land. Notably, aquatic realms feature important peculiarities: extraneous sounds from the air can be captured in so-called holo-soundscapes in freshwater and shallow coastal areas^61^, and particle motion (that accompanies sound) impacts aquatic organisms^62^. Advances in soundscape research are imminent as the freshwater acoustic research community is growing rapidly^63^. In the oceans, sound propagates comparatively far and multiple biomes can be sampled at once (e.g., recorders on the seafloor sample pelagic waters too) so that most ecosystems are covered. Still, coverage gaps may exist in rhodolith/Maërl beds. On land, sampling is biased towards biodiverse forests, and rocky habitats (young rocky pavements, lava flows and screes) but also some vegetated temperate ecosystems (cool temperate heathlands, temperate subhumid grasslands) are poorly sampled. Within the IUCN GET framework, soil soundscapes also belong to the terrestrial realm and to date only one spatio-temporally replicated dataset is in our database, despite recent studies^64,65^. Soil and benthic habitats may also be sampled with geophones to record infrasonic vibrations used to sense the environment (e.g., by insects, frogs, elephants, interstitial fauna and benthic fish)^66^.

Our database highlights well-known global sampling biases^67^ which could be resolved with collaboration and communication to remove cultural and socioeconomic barriers^68^. Technological progress for more affordable equipment renders PAM more accessible in lower-income countries^44,69^. However, high- and deep-sea work remains considerably more expensive, and tropical developing countries in particular often lack funding for marine programmes requiring large vessels, underwater vehicles, or cabled stations on the seafloor^46^. Our network currently consists of active members from 56 countries. Collaborative projects, shared sea missions, and equipment loans should promote the establishment of soundscape research communities^70^. However, equitable, collaborative efforts will also require capacity-building so that local researchers can independently process, analyse, and interpret PAM-derived data. Increased international collaboration with scientists and local stakeholders supporting citizen-science^71^ in heavily underrepresented regions would improve not only data coverage, but also representation and dialogue within the field.

Collaborative soundscape research relies on interoperable data. We harmonised metadata with a bottom-up approach leading to our global inventory. Comprehensive standards for PAM deployment, reporting, and data analysis are needed for enabling comparative, global analyses. However, such standards do not currently exist, even though initiatives for the marine realm are ongoing^72,73^. Few affordable solutions exist for sharing large audio data volumes^34^, underlining the need for distributed soundscape recording repositories^74^. Marine oil and gas industry projects routinely upload data as part of their efforts to mitigate noise impacts on marine animals^75^, but these recordings often focus on frequencies relevant to seismic prospecting and access may be restricted^76^. Furthermore, recording equipment requires calibration, sound detection spaces need to be measured^23,77^, and data privacy must be ensured on land^78^. In parallel, species sound libraries^42,79–81^ grow and continue to provide invaluable acoustic and taxonomic references without which soundscape analysis for soniferous organisms would be futile. International organisations such as the Global Biodiversity Information Facility (GBIF) will be key to roll-out standards (e.g., Darwin Core) in a top-down manner. For the moment, we encourage early planning of data archival. In the future, the Worldwide Soundscapes will interoperate with other databases to close remaining coverage gaps and enhance standardisation.

A unified approach to ecology with PAM is now possible. More comprehensive coverage is needed to decisively answer research, conservation, and management questions that PAM can address, so we encourage prospective contributors to curate their metadata and join our inclusive project. We show that a large portion of the PAM community is willing to collaborate across realms and form a global network. We advocate for a bolder PAM effort to inform the agenda of soundscape ecology^82^, reaching out to places where no sound has been recorded before as well as to urban settings^83^. The research community may open new avenues to study environmental effects on acoustic activity^56^, social species interactions^84^, human-wildlife relationships^40^, function and phylogeny^85^, soundscape effects on human health^86^, acoustic adaptation and niche hypotheses^87,88^, macroecological patterns across ecosystems^1^, and initiate an integrated approach to noise impacts on wildlife.

PAM is an established method that can be applied over large spatial and temporal scales. However, consistent, large-scale monitoring of the Earth’s soundscapes is direly needed and essential to establish baselines for historical trends^89^ and quantify rapid changes in biodiversity and natural systems. International funding schemes should integrate PAM into biodiversity monitoring platforms such as GBIF^90^ and GEO BON^91,92^. Soundscapes are just starting to be used in legislation as an ecosystem feature to be preserved^93^. Occupancy maps for soniferous wildlife obtained from PAM would underpin the evaluation of progress towards threat reduction and ecosystem service provision of the Kunming-Montreal Global Biodiversity Framework^94^. By building collaborations around the knowledge frontiers identified here, we can aim to comprehensively describe and understand the acoustic make-up of the planet.

## Acknowledgements

We are thankful to our colleagues who supported our work: Abigail Seybert, Abram B. Fleishman, Adriana C. Acero-Murcia, Adrien Charbonneau, Adrià López-Baucells, Aklavik, Alain Paquette, Albert García, Alexander C. Lees, Alexandra Buitrago-Cardona, Alfonso Zúñiga, Alfred Wegener Institute Helmholtz Centre for Polar and Marine Research, Allan G Oliveira, Almo Farina, Alpheous, Ambroise N. Zongo, Amy Oden, Ana Filipa Palmeirim, Ana Širović, Anamaria Dal Molin, Anastasia Viricheva, Andrea Gavio, Andrew N. Gweh, Andros T Gianuca, Annalea Beard, Anne Sourdril, Annebelle Kok, Anthony Tan, Anthony Truskinger, Anushka Rege, Aran Mooney, Arne Wenzel, Astrid Brekke Skrindo, Australian Institute of Marine Science, Base de Estudos do Pantanal - UFMS, Bastien Castagneyrol, Bea Maas, Benedictus Freeman, Bibiana Gómez-Valencia, Bob Dziak, Bobbi J. Estabrook, Borja Milá, Brian Miller, Bridget Maher, C. Lisa Mahon, Carolline Z Fieker, Carolyn Rosten, Cat Adshead, Catherine Potvin, Catrin Westphal, Chantal Huijbers, Charlie Griffiths, Charlotte Roemer, Chistophe Thébaud, Chong Chen, Christina Ieronymidou, Christine Cassin, Christine Erbe, Christopher Clark, Cicero Simao, Claire Attridge, Clara P. Amorim, Claudia A. Medina-Uribe, Claudia Ascencio-Elizondo, Clément Lemarchand, Colin Bates, Colin KC Wen, Conner Partaker, Cowichan Tribes Lulumexun department - Fisheries and Marine projects staff, Craig A Radford, Curtin University, Cássio Alencar Nunes, Cécile Albert, Cédric Gervaise, Dadang Dwi Putra, Daisy Dent, Dan Warren, Daniel Mihai Toma, Daniel Rieker, Danny Swainson, David Lecchini, David Tucker, David Watson, Dawit Yemane, Deanna Clement, Dedi Rahman, Denise Risch, Dennis Higgs, Dennis Kühnel, Dexter Hodder, Didier Casane, Diego Alejandro Gómez-Montes, Diego Balbuena, Diego Gil, Diego Pavón Jórdan, Diogo Provete, Dorgival Diogenes, Dustin Whalen, Edho Walesa Prabowo, Eduardo Rodríguez Martinez, Elisa Schutz, Elizabeth Clingham, Eliziane Oliveira, Ellen Mc Arthur, Elma Kay, Eloy Revilla, Emanuel Stefan Baltag, Emilia Sokolowska, Emilio Tan, En-Hao Liu, Enoc Martinez, Eric Parmentier, Eric Rakotomalala, Erich Fischer, Erika Berenguer, Estância Mimosa Ecoturismo, Evan Economo, Fabio De Leo Cabrera, Faisal Ali Awanrali Khan, Fang-Yee Lin, Fannie W. Shabangu, Fernando Hidalgo, Filipe F de Deus, Flávio Rodrigues, Frederic Sinniger, Frode Fossøy, Frédéric Bertucci, Frédéric Sèbe, Gabriel Leite, Gabriel leite, George Best, Gerard Bota, Germany, Giselle Mahung, Gonzalo Pérez-Rosales, Gonçalo Silva, Greg and Lisa Morgan, Groupe Chiroptères de Guyane, Guillermo McPherson, Hailey Davies, Hajar, Hannah Reyes, Hanneline Smit-Robinson, Harris Papadopoulos, Heriniaina Randrianarison, Hervé Jourdan, Hila shamon, Hiromi Kayama Watanabe, Holger Kreft, Héloïse Rouzé, Ian Avery Bick, Ilias Karmiris, Irfan Fitriawan, Isabelle Côté, Italo A A Rondon, Ivin Wilfred Raj, Jacques Keumo Kuenbou, Jakob Tougaard, Jamie Ratliff, Janice Ser Huay Lee, Jason Gedamke, Jasper Kanes, Jean Pierre Castro Namuche, Jean-Yves Royer, Jeff Robinson, Jennifer Clark, Jens-Georg Fischer, Jessica Deichmann, Jessica Qualley, Jodanne Pereira, Johan Diepstraten, Johanna Järnegren, John Ewen, John Hildebrand, John Ryan, Jos Barlow, Jose A. Hernandez-Blanco, Jose Leon, Jose Manuel Ochoa-Quintero, José Luis Mena, José Nilson Santos, José Wagner, João Gama Monteiro, Juan Diego Tovar, Juliana Carolina Herran García, Julie Perrot, Julio Baumgarten, Junior Tremblay, Just T. Bayle Sempere, Justine Magbanua, Jérémy Anso, Jérémy Froidevaux, Jérémy Froideveaux, Jérôme Sueur, Karen H. Beard, Katie Huong, Katie Innes, Kelsie Murchie, Ken Otter, Ken P. Findlay, Kenneth Dudley, Kevin Darras, Kevin G. Borja-Acosta, Kieran Cox, Kinga Buda, Kris Harmon, Laela Sayigh, Laetitia Hédouin, Laure Desutter-Grandcolas, Laurel Symes, Lauren Kuehne, Laurence H. De Clippele, Laurent Legendre, Leeann Henry, Leila Hatch, Lenka Brousset, Liana Chesini Rossi, Lin Schwarzkopf, Linilson Padovese, Lisa Loseto, Louise Emmerson, Lourdes Maigua Medrano, Luca Iacobucci, Lucia Di Iorio, Luciana Nacimento, Ludovic Crochard, Luis P. da Silva, Lény Lego, M. Clara P. Amorim, Malika René-Trouillefou, Mao-Ning Tuanmuu, Marc Fernandez, Marc Lammers, Marcia Maia, Marconi Campos, Marconi Campos Cerqueira, Marcos Vaira, Marcus Rowcliffe, Maria Cielo Bazterrica, Marilyn Beauchaud, Marina Puebla-Aparicio, Marlize Muller, Martim Melo, María Jesús García Bianco, Mathias Igulu, Matias Carandell, Matthew J. Weldy, Matthew K Pine, Matthew Slater, Mauricio Akmentins, Max Ritts, Michael Pashkevich, Michael Scherer-Lorenzen, Michael Towsey, Miguel Pessanha Pais, Mike van der Schaar, Milton Cezar Ribeiro, Mina Anders, Muhammad Justi Makmun Jusrin, Nadia Pieretti, Niaina Nirina Mahefa Andriamavosoloarisoa, Nicholas Brown, Nicholas Friedman, Nick Gardner, Nicolas Farrugia, Nicolas Hette-Tronquart, Nicolas Pajusco, Nikos Fakotakis, Noam Leader, Noelia Ríos, Nuno Faria, Ocean Acoustics Group, Ocean Networks Canada Society, Olaf Boebel, Omar Hiram Martinez Castillo, Paul McDonald, Paulo J. Fonseca, Pedro B. Lopes, Peter Ellis, Peter Taylor, Phillip Eichinski, Pooja Choksi, Pousada Xaraés, Pshemek Zdroik, Rainforest Connection (RFCx), Refúgio Ecológico Caiman, Refúgio da Ilha Ecolodge, Rex K. Andrew, Ricardo Duarte, Richard Fuller, Richard Kinsey, Rob Rempel, Robert John Young, Robert McCauley, Robert Spaan, Robert Young, Rodrigo Silva, Rouvah Andriafanomezantsoa, Saki Harii, Samuel Challéat, Samuel Haché, Sandra Mueller, Sara Gonçalves, Sarah Rowley, Sarika Khanwilkar, Sean Connell, Sean Martin, Sebastian Muñoz Duque, Shanghai Aquatic Wildlife Conservation and Research Center, Shannon MacPhee, Shinsuke Kawagucci, Sierra Hunicutt, Sierra Jarriel, Sigal Balshine, Silent cities consortium, Simon Childerhouse, Sina Weier, Sofie Van Parijs, Soledad Gaston, Stan Dosso, Stanislas Wroza, Stephanie Plön, Stephen Insley, Stuart Marsden, Stéphan Jacquet, Stéphane Père, Susanna Piovano, Sérgio Moreira, Tatiana Atemasova, Tetsuya Miwa, The Hunters and Trappers Committees of Inuvik, The Inuvialuit-Canada Fisheries Joint Management Committee, Thiago M Ventura, Thomas Sattler, Todor Ganchev, Tomas Altamirano, Tomasz Stanislaw Osiejuk, Tomaz Melo, Tomonari Akamatsu, Tristan Louth-Robins, Tulio Rossi, Tómas Grétar Gunnarsson, Under The Pole Consortium, Valentin Chevalier, Valerie Linden, Valéria Tavares, Vijay Ramesh, Vincent Médoc, Vu Dinh Thong, Vy Tram Nguyen, Wen-Ling Tsai, Will Duguid, Yang Liu, Yenifer Herrera-Varón, You-Fang Chen, Yue Qiu, Yuhang Song, Zehava Sigal, Zuania Colón-Piñeiro, Zuzana Burivalova, and Tuktoyaktuk, e, Çağlar Akçay, and Éric Parmentier.

