## Supplementary material for "Worldwide Soundscapes: a synthesis of passive acoustic monitoring across realms": Figure S1

### Supporting Information

***Table S1****: Summary statistics for the geographical distribution of sampling sites in the different realms, across latitudinal and topographical gradients*

| **realm** | **Mean latitude, range [°]** | **Mean topography, range [m]** |
| --- | --- | --- |
| Freshwater | 35 (-32,57) | 191 (0,3770) |
| Freshwater-Marine | 22 (-32,50) | -6 (-51,1) |
| Marine | 15 (-69,72) | -407 (-10900,110) |
| Marine-Freshwater-Terrestrial | 28 (-38,70) | 0 (-9,10) |
| Marine-Terrestrial | 33 (-16,56) | 65 (-20,450) |
| Subterranean | -6 (-6,-6) | 653 (277,810) |
| Subterranean-Freshwater | 22 (22,23) | NaN (Inf,-Inf) |
| Terrestrial | 37 (-69,70) | 637 (0,4548) |
| Terrestrial-Freshwater | 23 (-31,53) | 400 (-6,2024) |

***Table S2****: Spatial and temporal sampling coverage in terms of sampling sites and sampling days per site, within ecosystem functional groups, grouped by biomes within realms, following the IUCN Global Ecosystems Typology**^95^**, based on soundscape meta-datasets included in the Worldwide Soundscapes database. Some low temporal coverage values may be rounded to 0.*

| **Realm** | **Biome** | **Functional group** | **Area [km^2]** | **Sites** | **Temporal coverage per site [d]** |
| --- | --- | --- | --- | --- | --- |
| Terrestrial | Tropical-subtropical forests biome | Tropical/Subtropical lowland rainforests | 8343493 | 1350 | 34 |
| Terrestrial | Tropical-subtropical forests biome | Tropical/Subtropical montane rainforests | 1650275 | 245 | 95 |
| Terrestrial | Tropical-subtropical forests biome | Tropical/Subtropical dry forests and thickets | 5395719 | 138 | 236 |
| Terrestrial | Tropical-subtropical forests biome | Tropical heath forests | 223066 | 64 | 89 |
| Terrestrial | Temperate-boreal forests and woodlands biome | Deciduous temperate forests | 4737664 | 3172 | 18 |
| Terrestrial | Temperate-boreal forests and woodlands biome | Boreal and temperate high montane forests and woodlands | 12171277 | 2712 | 15 |
| Terrestrial | Temperate-boreal forests and woodlands biome | Warm temperate laurophyll forests | 2831170 | 224 | 1 |
| Terrestrial | Temperate-boreal forests and woodlands biome | Temperate pyric sclerophyll forests and woodlands | 172491 | 78 | 684 |
| Terrestrial | Temperate-boreal forests and woodlands biome | Oceanic cool temperate rainforests | 855944 | 36 | 161 |
| Terrestrial | Temperate-boreal forests and woodlands biome | Temperate pyric humid forests | 207743 | 7 | 1477 |
| Terrestrial | Shrublands and shrubby woodlands biome | Seasonally dry temperate heath and shrublands | 1820671 | 190 | 264 |
| Terrestrial | Shrublands and shrubby woodlands biome | Seasonally dry tropical shrublands | 246784 | 7 | 4 |
| Terrestrial | Shrublands and shrubby woodlands biome | Cool temperate heathlands | 190755 | 0 | 0 |
| Terrestrial | Shrublands and shrubby woodlands biome | Young rocky pavements, lava flows and screes | 1752367 | 0 | 0 |
| Terrestrial | Savannas and grasslands biome | Trophic savannas | 3759470 | 214 | 4 |
| Terrestrial | Savannas and grasslands biome | Pyric tussock savannas | 15732745 | 106 | 986 |
| Terrestrial | Savannas and grasslands biome | Temperate woodlands | 7216911 | 37 | 1056 |
| Terrestrial | Savannas and grasslands biome | Hummock savannas | 684073 | 10 | 365 |
| Terrestrial | Savannas and grasslands biome | Temperate subhumid grasslands | 9676075 | 0 | 0 |
| Terrestrial | Polar/alpine (cryogenic) biome | Polar tundra and deserts | 5528355 | 304 | 72 |
| Terrestrial | Polar/alpine (cryogenic) biome | Temperate alpine grasslands and shrublands | 4881364 | 287 | 26 |
| Terrestrial | Polar/alpine (cryogenic) biome | Tropical alpine grasslands and herbfields | 177047 | 58 | 141 |
| Terrestrial | Polar/alpine (cryogenic) biome | Polar/alpine cliffs, screes, outcrops and lava flows | 1806749 | 6 | 34 |
| Terrestrial | Polar/alpine (cryogenic) biome | Ice sheets, glaciers and perennial snowfields | 16715155 | 1 | 4 |
| Terrestrial | Intensive land-use biome | Plantations | 19445793 | 501 | 5 |
| Terrestrial | Intensive land-use biome | Urban and industrial ecosystems | 9498980 | 400 | 21 |
| Terrestrial | Intensive land-use biome | Annual croplands | 23597212 | 331 | 29 |
| Terrestrial | Intensive land-use biome | Derived semi-natural pastures and old fields | 22262354 | 296 | 139 |
| Terrestrial | Intensive land-use biome | Sown pastures and fields | 26764931 | 193 | 2 |
| Terrestrial | Deserts and semi-deserts biome | Sclerophyll hot deserts and semi-deserts | 3239850 | 32 | 1678 |
| Terrestrial | Deserts and semi-deserts biome | Semi-desert steppe | 13325240 | 23 | 1724 |
| Terrestrial | Deserts and semi-deserts biome | Succulent or Thorny deserts and semi-deserts | 1836801 | 15 | 1 |
| Terrestrial | Deserts and semi-deserts biome | Hyper-arid deserts | 4590239 | 6 | 32 |
| Terrestrial | Deserts and semi-deserts biome | Cool deserts and semi-deserts | 7399041 | 1 | 2 |
| Marine | Pelagic ocean waters biome | Epipelagic ocean waters | 360419256 | 180 | 506 |
| Marine | Pelagic ocean waters biome | Mesopelagic ocean water | 334825581 | 21 | 828 |
| Marine | Pelagic ocean waters biome | Sea ice | 34040730 | 20 | 326 |
| Marine | Pelagic ocean waters biome | Bathypelagic ocean waters | 319407004 | 15 | 687 |
| Marine | Pelagic ocean waters biome | Abyssopelagic ocean waters | 272382441 | 1 | 169 |
| Marine | Marine shelf biome | Photic coral reefs | 5418204 | 187 | 43 |
| Marine | Marine shelf biome | Subtidal mud plains | 31654790 | 81 | 440 |
| Marine | Marine shelf biome | Subtidal sand beds | 31654790 | 53 | 329 |
| Marine | Marine shelf biome | Subtidal rocky reefs | 31654790 | 37 | 240 |
| Marine | Marine shelf biome | Seagrass meadows | 3054851 | 27 | 238 |
| Marine | Marine shelf biome | Kelp forests | 2409185 | 9 | 13 |
| Marine | Marine shelf biome | Shellfish beds and reefs | 6944349 | 4 | 505 |
| Marine | Marine shelf biome | Photo-limited marine animal forests | 31276008 | 2 | 2 |
| Marine | Marine shelf biome | Upwelling zones | 3052042 | 1 | 95 |
| Marine | Marine shelf biome | Rhodolith/Maërl beds |  | 0 | 0 |
| Marine | Deep sea floors biome | Continental and island slopes | 19555151 | 42 | 453 |
| Marine | Deep sea floors biome | Chemosynthetic-based-ecosystems (CBE) | 1716366 | 8 | 328 |
| Marine | Deep sea floors biome | Deepwater biogenic beds | 203723553 | 7 | 115 |
| Marine | Deep sea floors biome | Abyssal plains | 247963714 | 5 | 1048 |
| Marine | Deep sea floors biome | Submarine canyons | 4386833 | 4 | 341 |
| Marine | Deep sea floors biome | Hadal trenches and troughs | 6955196 | 1 | 359 |
| Marine | Deep sea floors biome | Seamounts, ridges and plateaus | 57050685 | 1 | 2 |
| Marine | Anthropogenic marine biome | Submerged artificial structures | 8762580 | 10 | 182 |
| Marine | Anthropogenic marine biome | Marine aquafarms | 9343092 | 6 | 21 |
| Freshwater | Rivers and streams biome | Permanent lowland rivers | 1351261 | 80 | 89 |
| Freshwater | Rivers and streams biome | Seasonal lowland rivers | 1304042 | 15 | 2 |
| Freshwater | Rivers and streams biome | Large lowland rivers | 718843 | 11 | 8 |
| Freshwater | Rivers and streams biome | Episodic arid rivers | 2136890 | 1 | 13 |
| Freshwater | Rivers and streams biome | Freeze-thaw rivers and streams | 13072135 | 1 | 378 |
| Freshwater | Rivers and streams biome | Seasonal upland streams | 16213367 | 1 | 19 |
| Freshwater | Rivers and streams biome | Permanent upland streams | 7085982 | 0 | 0 |
| Freshwater | Lakes biome | Small permanent freshwater lakes | 166635 | 93 | 207 |
| Freshwater | Lakes biome | Large permanent freshwater lakes | 643901 | 39 | 346 |
| Freshwater | Lakes biome | Permanent salt and soda lakes | 428359 | 1 | 8 |
| Freshwater | Lakes biome | Artesian springs and oases | 475490 | 0 | 0 |
| Freshwater | Lakes biome | Ephemeral freshwater lakes | 73255 | 0 | 0 |
| Freshwater | Lakes biome | Ephemeral salt lakes | 1136371 | 0 | 0 |
| Freshwater | Lakes biome | Freeze-thaw freshwater lakes | 12099415 | 0 | 0 |
| Freshwater | Lakes biome | Geothermal pools and wetlands | 528692 | 0 | 0 |
| Freshwater | Lakes biome | Seasonal freshwater lakes | 454622 | 0 | 0 |
| Freshwater | Lakes biome | Subglacial lakes | 70903 | 0 | 0 |
| Freshwater | Artificial wetlands biome | Rice paddies | 5791268 | 18 | 16 |
| Freshwater | Artificial wetlands biome | Constructed lacustrine wetlands | 37224442 | 11 | 4 |
| Freshwater | Artificial wetlands biome | Canals, ditches and drains | 13216648 | 3 | 131 |
| Freshwater | Artificial wetlands biome | Large reservoirs | 8726072 | 1 | 873 |
| Freshwater | Artificial wetlands biome | Freshwater aquafarms | 20607575 | 0 | 0 |
| Subterranean | Subterranean lithic biome | Aerobic caves | 12688679 | 7 | 1 |
| Subterranean | Subterranean lithic biome | Endolithic systems | 510065622 | 0 | 0 |
| Subterranean | Anthropogenic subterranean voids biome | Anthropogenic subterranean voids | 18126519 | 0 | 0 |
| Marine-Terrestrial | Supralittoral coastal biome | Large seabird and pinniped colonies |  | 14 | 3 |
| Marine-Terrestrial | Supralittoral coastal biome | Coastal shrublands and grasslands | 1770542 | 13 | 6 |
| Marine-Terrestrial | Shorelines biome | Sandy Shorelines | 1089123 | 7 | 6 |
| Marine-Terrestrial | Shorelines biome | Boulder and cobble shores | 728035 | 0 | 0 |
| Marine-Terrestrial | Shorelines biome | Muddy Shorelines | 145654 | 0 | 0 |
| Marine-Terrestrial | Shorelines biome | Rocky Shorelines | 1452282 | 0 | 0 |
| Marine-Terrestrial | Anthropogenic shorelines biome | Artificial shorelines | 681154 | 0 | 0 |
| Subterranean-Freshwater | Subterranean freshwaters biome | Underground streams and pools | 12688679 | 6 | 86 |
| Subterranean-Freshwater | Subterranean freshwaters biome | Groundwater ecosystems | 17416482 | 0 | 0 |
| Subterranean-Freshwater | Anthropogenic subterranean freshwaters biome | Flooded mines and other voids | 16782429 | 0 | 0 |
| Subterranean-Freshwater | Anthropogenic subterranean freshwaters biome | Water pipes and subterranean canals | 32101390 | 0 | 0 |
| Terrestrial-Freshwater | Palustrine wetlands biome | Seasonal floodplain marshes | 1220146 | 86 | 120 |
| Terrestrial-Freshwater | Palustrine wetlands biome | Tropical flooded forests and peat forests | 1239398 | 79 | 25 |
| Terrestrial-Freshwater | Palustrine wetlands biome | Boreal, temperate and montane peat bogs | 13471527 | 56 | 10 |
| Terrestrial-Freshwater | Palustrine wetlands biome | Subtropical/temperate forested wetlands | 22557222 | 16 | 208 |
| Terrestrial-Freshwater | Palustrine wetlands biome | Permanent marshes | 397429 | 12 | 184 |
| Terrestrial-Freshwater | Palustrine wetlands biome | Boreal and temperate fens | 15012885 | 8 | 15 |
| Terrestrial-Freshwater | Palustrine wetlands biome | Episodic arid floodplains | 467656 | 0 | 0 |
| Subterranean-Marine | Subterranean tidal biome | Anchialine caves | 93142 | 0 | 0 |
| Subterranean-Marine | Subterranean tidal biome | Anchialine pools | 93142 | 0 | 0 |
| Subterranean-Marine | Subterranean tidal biome | Sea caves | 1429873 | 0 | 0 |
| Marine-Freshwater-Terrestrial | Brackish tidal biome | Coastal river deltas | 652553 | 64 | 111 |
| Marine-Freshwater-Terrestrial | Brackish tidal biome | Coastal saltmarshes and reedbeds | 68921 | 16 | 67 |
| Marine-Freshwater-Terrestrial | Brackish tidal biome | Intertidal forests and shrublands | 150345 | 2 | 438 |
| Freshwater-Marine | Semi-confined transitional waters biome | Permanently open riverine estuaries and bays | 3184031 | 52 | 125 |
| Freshwater-Marine | Semi-confined transitional waters biome | Intermittently closed and open lakes and lagoons | 1431473 | 10 | 21 |
| Freshwater-Marine | Semi-confined transitional waters biome | Deepwater coastal inlets | 264394 | 0 | 0 |

***Table S3****: Summary statistics of the soundscape case studies*

| **Functional group** | **Diel time windows** | **Sites** | **Dates** | **Biophony occupancy** | **Geophony occupancy** | **Anthropophony occupancy** | **Silence occupancy** |
| --- | --- | --- | --- | --- | --- | --- | --- |
| Large lowland rivers  * | evening, morning, night, noon | 4 | 2014-06-20 - 2014-06-21 | 0.22 | 0 | 0 | 0.78 |
| Riverine estuaries | evening, morning, night, noon | 3 | 2014-01-12 - 2015-05-02 | 0.49 | 0 | 0.14 | 0.42 |
| Tropical montane rainforests | evening, morning, night, noon | 4 | 2021-05-04 - 2021-05-05 | 0.22 | 0.05 | 0.01 | 0.72 |
| Island slopes | noon | 4 | 2019-07-25 - 2019-08-05 | 0 | 0 | 0.32 | 0.68 |
| Photic coral reefs | evening, morning, night, noon | 3 | 2017-06-06 | 0.75 | 0.02 | 0.05 | 0.22 |
| Plantations | evening, morning, night, noon | 4 | 2016-06-15 - 2016-06-16 | 0.29 | 0 | 0.05 | 0.67 |
| Tropical lowland rainforests | evening, morning, night, noon | 4 | 2016-06-15 - 2016-06-16 | 0.61 | 0 | 0 | 0.39 |
| Bathypelagic ocean waters | evening, morning, night, noon | 3 | 2022-03-21 - 2022-03-22 | 0.03 | 0 | 0 | 0.96 |
| Small permanent freshwater lakes | evening, morning, night, noon | 3 | 2023-03-08 | 0.17 | 0.03 | 0 | 0.62 |
| Urban ecosystems  + | evening, morning, night, noon | 8 | 2022-02-24 - 2023-05-27 | 0.16 | 0.05 | 0.43 | 0.42 |
| Aerobic caves  ** | evening, morning, night | 4 | 2023-06-05 - 2023-06-14 | 0.26 | 0 | 0 | 0.71 |
| Polar outcrops  # | evening, morning | 4 | 2021-12-23 - 2021-12-24 | 0.02 | 0.14 | 0.04 | 0.82 |

*: 5-minute recordings recomposed from five 1-min recordings recorded over 1 hour with a 1/10 duty-cycle.
**: 5-minute recordings recomposed from 5-second recordings recorded over half an hour with a 5 s / 25 s duty cycle
#: 24 kHz sampling frequency, no noon or night at this latitude
+: human voices removed with high-pass filter
~: closest 10 min to solar times chosen from 30/60 min duty-cycled recordings in Montreal (maximum offset: 10 min)

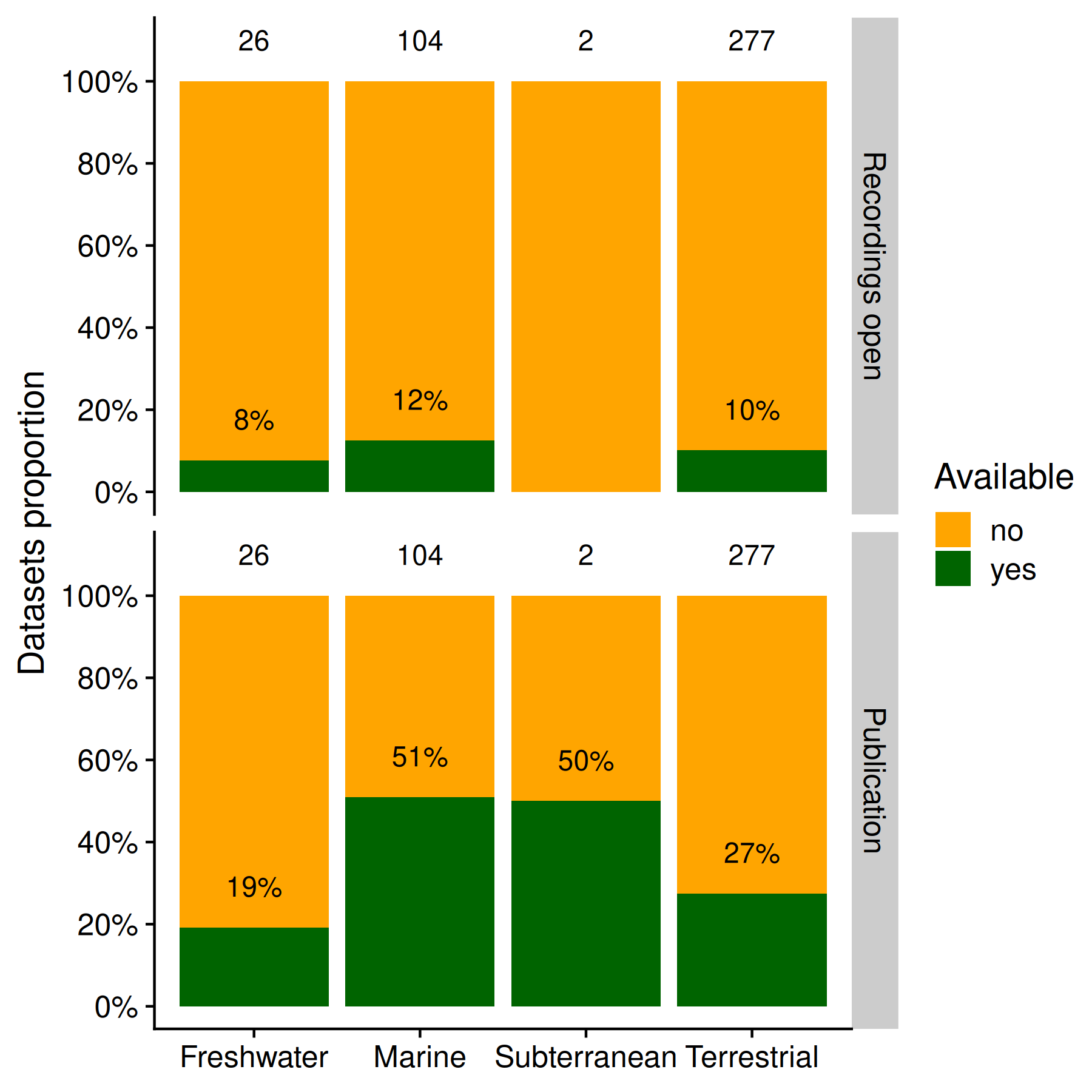

***Figure S1****: A) Availability of recordings and published articles of the datasets in the different core realms, displayed as percentages, as determined by the presence of openly-accessible internet links and DOIs. Total dataset number indicated above bars.*

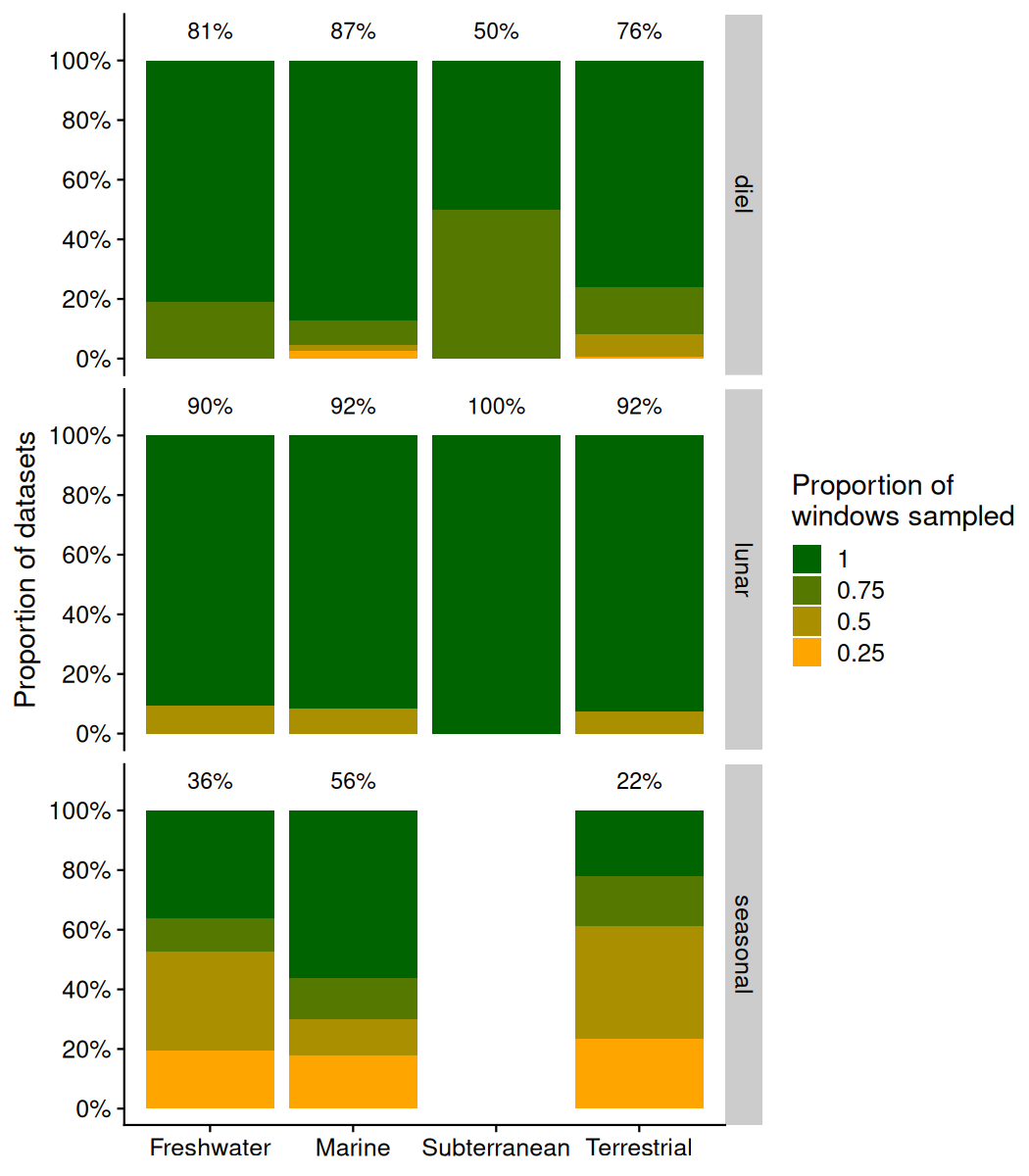

***Figure S2****: Sampling completeness as measured by the proportion of different time windows covered by acoustic recordings out of the total number of time windows, for each time cycle and realm. Percentages above stacked bars indicate the percentage of datasets that sample all time windows of each cycle.*

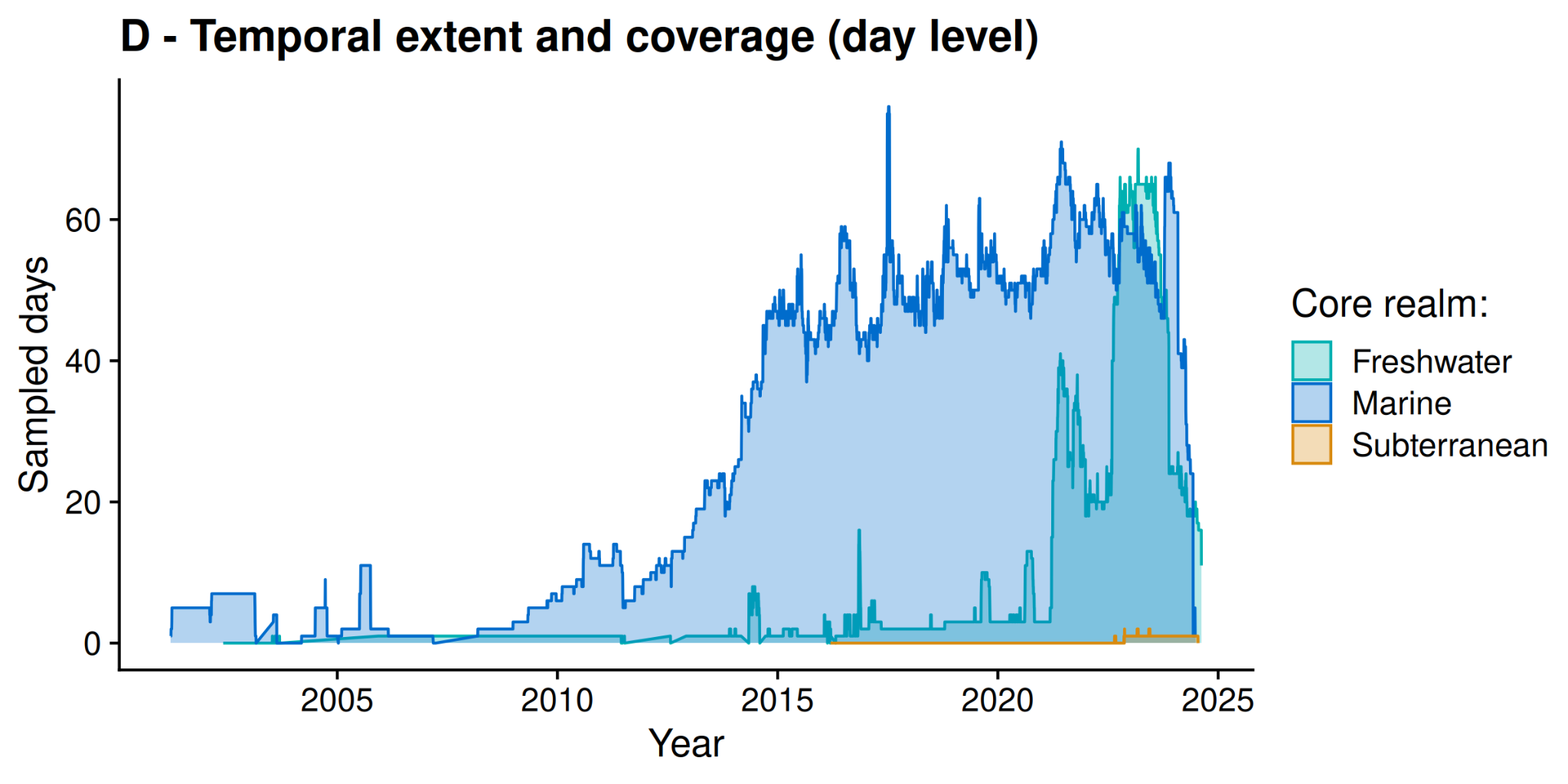

***Figure S3****: Timeline of temporal coverage across realms, without terrestrial sites*

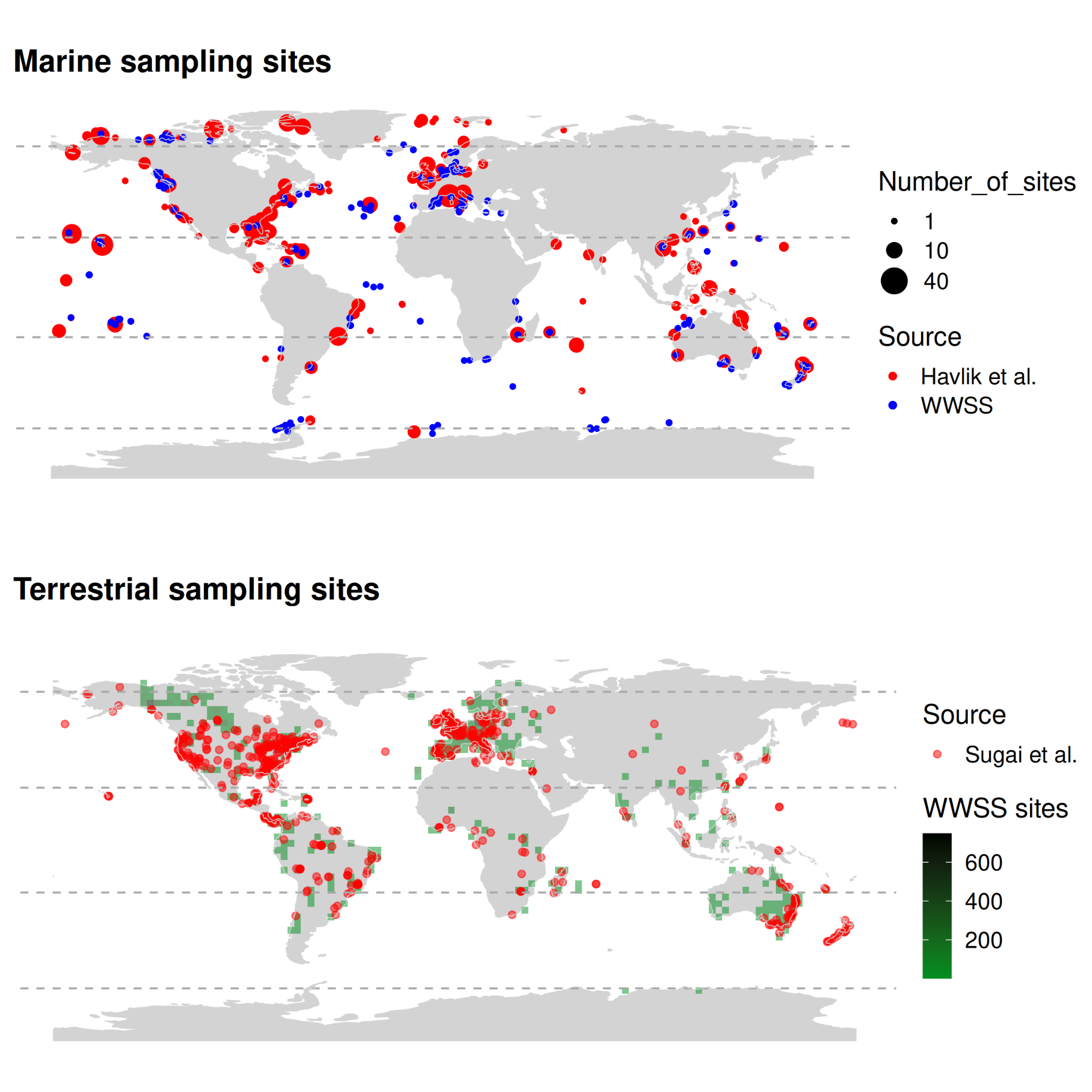

***Figure S4****: A) Geographic locations of the Worldwide Soundscapes (WWSS) database marine sampling sites and of Havlik et al.'s**^15^* *datasets. B) Geographic locations of the Worldwide Soundscapes (WWSS) database terrestrial sampling sites and of Sugai et al.'s**^14^* *sampling sites (reproduced with permission).*
